# The AEGEAN-169 clade of bacterioplankton is synonymous with SAR11 subclade V (HIMB59) and metabolically distinct

**DOI:** 10.1101/2023.02.22.529538

**Authors:** Eric W. Getz, V. Celeste Lanclos, Conner Y. Kojima, Chuankai Cheng, Michael W. Henson, Max Emil Schön, Thijs J. G. Ettema, Brant C. Faircloth, J. Cameron Thrash

## Abstract

Bacterioplankton of the SAR11 clade are the most abundant marine microorganisms and consist of numerous subclades spanning Order level divergence (*Pelagibacterales*). The assignment of the earliest diverging subclade V (a.k.a. HIMB59) to the *Pelagibacterales* is highly controversial, with multiple recent phylogenetic studies placing them completely separate from SAR11. Other than through phylogenomics, subclade V has not received detailed examination due to limited genomes from this group. Here, we assessed the ecogenomic characteristics of subclade V to better understand the role of this group in comparison to the *Pelagibacterales*. We used a new isolate genome, recently released single amplified genomes (SAGs) and metagenome-assembled genomes (MAGs), and previously established SAR11 genomes to perform a comprehensive comparative genomics analysis. We paired this analysis with recruitment of metagenomes spanning open ocean, coastal, and brackish systems. Phylogenomics, average amino acid identity, and 16S rRNA gene phylogeny indicate that SAR11 subclade V is synonymous with the ubiquitous AEGEAN-169 clade, and support the contention that this group represents a taxonomic Family. AEGEAN-169 shared many bulk genome qualities with SAR11, such as streamlining and low GC content, but genomes were generally larger. AEGEAN-169 had overlapping distributions with SAR11 but was metabolically distinct from SAR11 in its potential to transport and utilize a broader range of sugars as well as in transport of trace metals and thiamin. Thus, regardless of the ultimate phylogenetic placement of AEGEAN-169, these organisms have distinct metabolic capacities that likely allow them to differentiate their niche from canonical SAR11 taxa.

**IMPORTANCE:** One goal of marine microbiologists is to uncover the roles various microorganisms are playing in biogeochemical cycles. Success in this endeavor relies on differentiating groups of microbes and circumscribing their relationships. An early-diverging group (subclade V) of the most abundant bacterioplankton, SAR11, has recently been proposed as a separate lineage that does not share a most recent common ancestor. But beyond phylogenetics, little has been done to evaluate how these organisms compare with SAR11. Our work leverages dozens of new genomes to demonstrate the similarities and differences between subclade V and SAR11. In our analysis, we also establish that subclade V is synonymous with a group of bacteria established from 16S rRNA gene sequences, AEGEAN-169. Subclade V/AEGEAN-169 has clear metabolic distinctions from SAR11 and their shared traits point to remarkable convergent evolution if they do not share a most recent common ancestor.

## INTRODUCTION

SAR11 are aerobic chemoorganoheterotrophs that comprise the largest fraction of bacterioplankton in the global ocean (1). Hallmarks of the group include streamlined genomes with high coding densities and few pseudogenes or gene duplications (2–4); unique requirements for amino acids, osmolytes, and C1 compounds (1); and a paucity of canonical regulatory suites (4). Five major SAR11 subclades have been classified and defined through ecogenomic observations during the preceding decades using 16S rRNA gene phylogenetic and whole genome phylogenomic approaches (1, 5–7). SAR11 is currently classified as a taxonomic order (*Pelagibacterales*), and the subclades represent genus to family level distinctions. The majority of SAR11 subclades are found in the epipelagic region, with the predominant subclade being Ia (7), however, subclades Ic and IIb can be found within the mesopelagic and bathypelagic (7–10). Surface water genomes have an average size of 1.33 Mbp, contrasting with that of deeper water genomes that average 1.49 Mbp (4, 10). The earliest diverging subclade V comprises two groups – Va shares a surface summer distribution with Ia in the Sargasso Sea, whereas Vb has both a surface and sub-euphotic distribution (7).

Although a stable member of SAR11 in rRNA gene phylogenies (7, 11), the inclusion of subclade V within SAR11 has recently been questioned by advanced phylogenomic approaches using new data (12, 13). Initially, some of these results were questionable due to the availability of only a single genome (HIMB59 - (3)) representing subclade V. However, reconstruction of subclade V MAGs provided additional genomic signal and the use of methods to correct for compositional biases placed HIMB59-type organisms on a separate branch of the Alphaproteobacteria (13, 14). Nevertheless, analyses of the HIMB59 genome indicated numerous similarities with SAR11, including the small size, low GC content, and conservation of similar metabolic pathways (3). Based on the genomic and ecological similarities with SAR11, deeper investigation of HIMB59-type organisms is warranted to understand their convergence with SAR11.

Early studies with 16S rRNA gene cloning have also defined a sister group to SAR11 that was given the name AEGEAN-169 (15). The group has a cosmopolitan distribution, identified in many regions including the Xiamen Sea, the San Pedro Ocean Time Series (SPOT), the South Pacific Gyre, and the Adriatic Sea (16–20). AEGEAN-169 was especially abundant in surface waters of the South Pacific Gyre and the Sargasso Sea, where numerous single-cell genomes were recently obtained, supporting the hypothesis of an ultraoligotrophic lifestyle (18, 19, 21). However, these organisms also respond to phytoplankton blooms (16), and AEGEAN-169 has also been observed at depths of 500 m or below at SPOT (17) and at 400 m in the NE Atlantic (22), as well as in coastal (23, 24) and reef (25) habitats. Seasonal blooms of AEGEAN-169 have been identified in the Mediterranean and Xiamen Seas through CARD-FISH and sequencing methodologies where their abundance was related to elevated CO2 concentrations and temperature increases (19, 26). As a result, AEGEAN-169 may play a key role in expanding and warming oligotrophic conditions, globally. AEGEAN-169 have also been implicated in phosphonate consumption (27), implicating another adaptation for the oligotrophic lifestyle.

While our knowledge of this group has improved, the AEGEAN-169 clade has not been examined thoroughly with comparative genomics, nor has its relationship to the SAR11 clade been formally established using modern phylogenomic techniques. AEGEAN-169 have sometimes been classified as belonging to the *Rhodospirillales* (16, 24, 26) and, more recently, were used as an outgroup to SAR11 within the Alphaproteobacteria (21, 27). Here, we present evidence from 16S rRNA gene phylogenetics and phylogenomics that AEGEAN-169 is a heterotopic synonym with SAR11 subclade V, also known as the HIMB59-type clade after the first isolate from the group (3). We do not attempt to reclassify the phylogeny of these organisms, as the close relationship between subclade V/HIMB59 and SAR11 has been examined in detail with advanced phylogenetic methods and appears to result from compositional artifacts (13, 14). Rather, we performed an extensive comparative genomics analysis using publicly available MAGs and SAGs from multiple databases (21, 28–30). We also include a closed genome from the second reported culture of this group, strain LSUCC0245, previously classified as a close relative to HIMB59 (31), and we provide the first physiological data for the clade resulting from this isolate. We aimed to define the taxonomy, distribution, and metabolic potential of AEGEAN-169/SAR11 subclade V/HIMB59 to better characterize its relationship to SAR11 *sensu stricto*.

## MATERIALS AND METHODS

### Genome sequencing and assembly of LSUCC0245

We previously isolated a close relative of HIMB59, strain LSUCC0245 (32). Due to the low densities of LSUCC0245 (mid-10^5^ cells ml^-1^), and an inability of this organism to grow in large volumes, sixty 50 ml cultures (Supplemental Information -https://doi.org/10.6084/m9.figshare.22027763) grown in JW2 (32) were aggregated to achieve sufficient volumes for DNA sequencing. Samples were harvested via 0.2 µm filtration (polycarbonate, Millipore) in late log phase. DNA was extracted using the Mobio PowerWater kit (Qiagen) with a 50 ml elution in water, and library preparation and sequencing were performed as described (33). Illumina HiSeq sequencing generated 1,925,078 paired-end, 150 bp reads. Genome assembly was performed as described (33). Briefly, reads were trimmed with Trimmomatic v0.38 (34), assembled with SPAdes v3.10.1 (35), and quality checked using Pilon v1.22 (36) after mapping reads to the assembly using BWA 0.7.17 (37). The assembly resulted in a single, circular contig, which was manually rotated approximately halfway between the original overlapping ends. Pilon was run on both the original contig and the rotated contig and detected no issues. Final coverage was 242x. The genome was annotated at IMG (https://img.jgi.doe.gov/) (38).

### Taxon selection

We used SAR11 genomes collected previously from GTDB (30, 33) and AEGEAN-169 genomes from the IMG database, the GORG-TROPICS SAGs database, the Microbiomics database, and the OceanDNA MAG catalog (21, 28, 29, 39). We initially used the HIMB59 and LSUCC0245 genomes, as well as AEGEAN-169 SAGs from GORG-TROPICS and our SAR11 genome collection as a starting dataset, and used FastANI v1.33 (40) with default settings to identify additional SAR11 and AEGEAN-169 genomes from the Microbiomics and the OceanDNA MAG datasets. We dereplicated our initial dataset of 814 genomes with dREP v3.4.0 (41) using *“dereplicate”* with default settings to produce a final dataset of 438 representatives including AEGEAN-169 and the SAR11 clade (Supplemental Information -https://doi.org/10.6084/m9.figshare.22027763).

### 16S rRNA gene phylogeny

We used barrnap v0.9 (42) to parse all available 16S rRNA genes from the 438 genomes and combined them with relevant AEGEAN-169 16S rRNA gene clones (15), four rRNA gene clones that had been previously classified as SAR11 subclades Va and Vb (7), and other Alphaproteobacteria (https://doi.org/10.6084/m9.figshare.22027763). We aligned extracted gene sequences with Muscle v3.8.1551 (43) using default settings and constructed the tree using IQ-Tree2 v3.8.1551 (44) using *“-b”* for traditional bootstrapping (n=100), and which selected the GTR+F+I+G4 model. The tree was visualized and formatted using iTOL v5 (45).

### 16S rRNA gene identity

To calculate 16S rRNA gene identity, we constructed a BLAST (46) database of the 16S rRNA gene sequences from SAR11 and AEGEAN-169 using makeblastdb v2.9.0 with database *“-type nucl”*. We then ran blastn v2.9.0 with *“-perc_identity 40”* and an e-value threshold of 1e-15 using the same 16S rRNA gene sequences to generate all pairwise 16S rRNA gene identities.

### Genome metrics

We calculated genome metrics for all genomes in the final dataset with CheckM v1.1.3 lineage_wf (47). We ran “checkm tree_qa” followed by “checkm lineage_set”. Continuing we ran “chekm analyze” followed by “checkm qa”. Relevant data including genome size, GC content, coding density, genome contamination, and genome completeness resulted from the check output. Estimated genome size was calculated using CheckM metrics as follows:

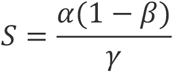

where *α* is the number of actual genome base pairs, *β* is predicted contamination, and *γ* is estimated completeness, as described previously (48).

### Pangenome construction and metabolic profiling

Pangenomic analyses were completed with Anvi’o v7.1 (49). First, we generated Anvi’o contigs databases using *“anvi-gen-contigs-database”*. We then ran a series of annotations, calling the contigs database. For Pfam (50) annotations, we ran *“anvi-run-pfams”*. For NCBI clusters of orthologous groups (COGs) (51) we ran *“anvi-run-ncbi-cogs”*. To import KEGG (52) annotations, we exported all amino acid sequences from respective contigs databases applying *“anvi-get-sequences-for-gene-calls”*. Amino acid sequences were input into the Ghostkoala (53) web application at KEGG (https://www.kegg.jp/ghostkoala/). Ghoastkoala output was parsed to match respective contigs databases and prepped using *“KEGG-to-anvio”*. To import KEGG functions, we employed *“anvi-import-functions”*. To generate a genomes database from the annotated contigs databases we used *“anvi-get-genomes-storage”*. Having generated a viable genomes database, we then employed *“anvi-pan-genome”* with a minbit setting of 0.5 and mcl-inflation set at 2 to construct a pangenome database. To identify enriched functions by subclade we affixed subclade metadata to the pangenome database using *“anvi-import-misc-data”*. Following this, we ran *“anvi-get-enriched-functions-per-pan-group”* calling COG_category, COG_function, KeggGhostkoala, and Pfam, respectively (54). A pangenome summary was exported via *“anvi-summarize”* (55, 56). The pangenome summary is available in Supplemental Information (https://doi.org/10.6084/m9.figshare.22027763).

### Phylogenomics

Genomes from AEGEAN-169 and SAR11 clade members were used for phylogenomics with conserved single-copy protein sequences as described previously (57). Briefly, single-copy orthologs were selected from the Anvi’o pangenomics output and all amino acid sequences were aligned and trimmed using Muscle v3.8.1551 and Trimal v1.4.1 with the “-automated1” flag, (43, 58). The individual alignments were concatenated using the geneStitcher.py script from the Utensils package (https://github.com/ballesterus/Utensils) (59) and the phylogeny was inferred from the unpartitioned, concatenated alignment (https://doi.org/10.6084/m9.figshare.22027763) using IQ-Tree2 v2.0.6 (44), that selected the best-fitting site rate substitution model (LG+F+R10), and *“-bb”* for ultrafast bootstrapping. The tree was visualized and formatted using iTOL v5 (45).

### Proteorhodopsin phylogenetics

To more accurately classify proteorhodopsin diversity across the different predicted variants, orthologous clusters from the Anvi’o pangenomics workflow that were annotated as rhodopsin proteins were aligned with reference sequences provided by Oded Beja (personal communication) using Muscle v3.8.1551, culled with Trimal v1.4.1 with the “-automated1” flag, and the phylogeny was inferred using IQ-Tree2 v2.0.6 (44), that selected the best-fitting site rate substitution model (VT+F+G4), and *“-bb”* for ultrafast bootstrapping. The tree was visualized and formatted using iTOL v5 (45). Proteorhodopsin tuning was assigned as previously described (60). The starting fasta file is available in Supplemental Information (https://doi.org/10.6084/m9.figshare.22027763).

### Metagenomic recruitment

Metagenomic samples were compiled from the following datasets: TARA Oceans, BIOGEOTRACES, MALASPINA, the Bermuda Atlantic Time Series (BATS), the Chesapeake, Delaware, and San Francisco Bays, the Hawaiian Ocean Time series (HOT), the Columbia River and Yaquina Bay, the Baltic Sea, Pearl River, Sapelo Island, Southern California Bight, and the northern Gulf of Mexico (61–69). We recruited reads from all datasets to the AEGEAN-169 genomes *via* RRAP (70–72). Post-recruitment, we assessed subclade distribution by summing all Reads Per Kilobases of genome per Million bases of metagenome sequence (RPKM) values for the genomes within each subclade and plotting them by depth, temperature, and salinity.

### HOT analysis

To assess seasonal distributions of AEGEAN-169, we used data from the Hawaiian Ocean Time series (HOT) that contained monthly samples for several different years. We sorted our global recruitment data to parse HOT-specific samples from Station ALOHA for the years 2004-2016. We then summed RPKM values respective to each subclade. We used Ocean Data Visualization (ODV) to sort summed RPKM data by subclade, month, and depth to interpret seasonality over a 12-month timeline (73).

### Growth experiments

LSUCC0245 was experimentally tested for growth ranges and optima as described previously (32). Briefly, we created artificial seawater media of different salinities through proportional dilution of the major salts. For the temperature-specific experiments, we used the isolation medium, JW2. Growth was measured with flow cytometry as described (32, 74).

### Data visualization

All custom scripts and underlying data for generating the metagenomic recruitment, 16S identity, and comparative metabolism figures are available in Supplemental Information (https://doi.org/10.6084/m9.figshare.22027763).

## RESULTS

### Genome reconstruction of LSUCC0245

Strain LSUCC0245 was isolated as previously reported from surface water near the Calcasieu Ship Channel jetties in Cameron, Louisiana and found to be most similar to HIMB59 based on 16S rRNA gene sequence similarity (31). The two organisms share 99.77% 16S rRNA gene identity. We recovered a complete, circularized genome for strain LSUCC0245 that was 1,493,989 bp with a 32.54% GC content and 1,585 predicted coding genes.

### Phylogenetics and taxonomy

We constructed a 16S rRNA gene tree using all recovered genes from the MAGs, SAGs, and isolates, as well as clones from the original AEGEAN-169 sequence report (15), using SAR11 and other Alphaproteobacteria as outgroups. We also included the subclade Va and Vb sequences previously used to delineate subclade V in SAR11 (7). We found that the Va and Vb sequences corresponded to two monophyletic groups containing all the 16S rRNA gene sequences from our genomes (including HIMB59 and LSUCC0245), as well as the AEGEAN-169 clone library sequences (**Fig. S1**). This topology demonstrates that the previously designated SAR11 subclade V is synonymous with AEGEAN-169, and we refer to the group by the latter name hereafter. AEGEAN-169 subclade I showed slightly deeper vertical branching in comparison to AEGEAN-169 subclade II.

The average 16S rRNA gene identity between AEGEAN-169 and SAR11 was 84.3% (**Fig. S2**). This indicates a likely Family level difference, but is near the boundary specification for Order classification at 82% (75). AEGEAN-169 within subclade I and II gene identities averaged 97.9% and 95.7%, indicating that each subclade corresponds to a species rank. Thus, the formerly defined Va and Vb groupings correspond to two distinct species groups within AEGEAN-169, and the entire clade likely represents at least a distinct Family.

To investigate the branching pattern between AEGEAN-169 subclades I and II, as well as within each subclade, we also constructed a phylogenomic tree of AEGEAN-169 and SAR11 using orthologous protein sequences extracted from the 438 genomes. The final translated alignment contained 28,837 amino acid positions. The monophyletic grouping of SAR11 and AEGEAN-169 can arise from compositional artifacts (13, 14), and we made no attempt to correct for these artifacts here. Rather, we only used SAR11 as an outgroup based on rRNA gene relationships (7, 11) (**Fig. S1**). Similarly to the 16S rRNA gene tree, we observed two distinct monophyletic subclades encompassing all AEGEAN-169 genomes wherein strain LSUCC0245 was sister to HIMB059 (**Fig. 1**). AEGEAN-169 subclade I was characterized by four distinct subgroups (Ia-Id) and subclade II was characterized by seven subgroups (IIa-IIg) defined through branching patterns. LSUCC0245 and HIMB59 were members of subgroup Ib.

**Figure 1.**
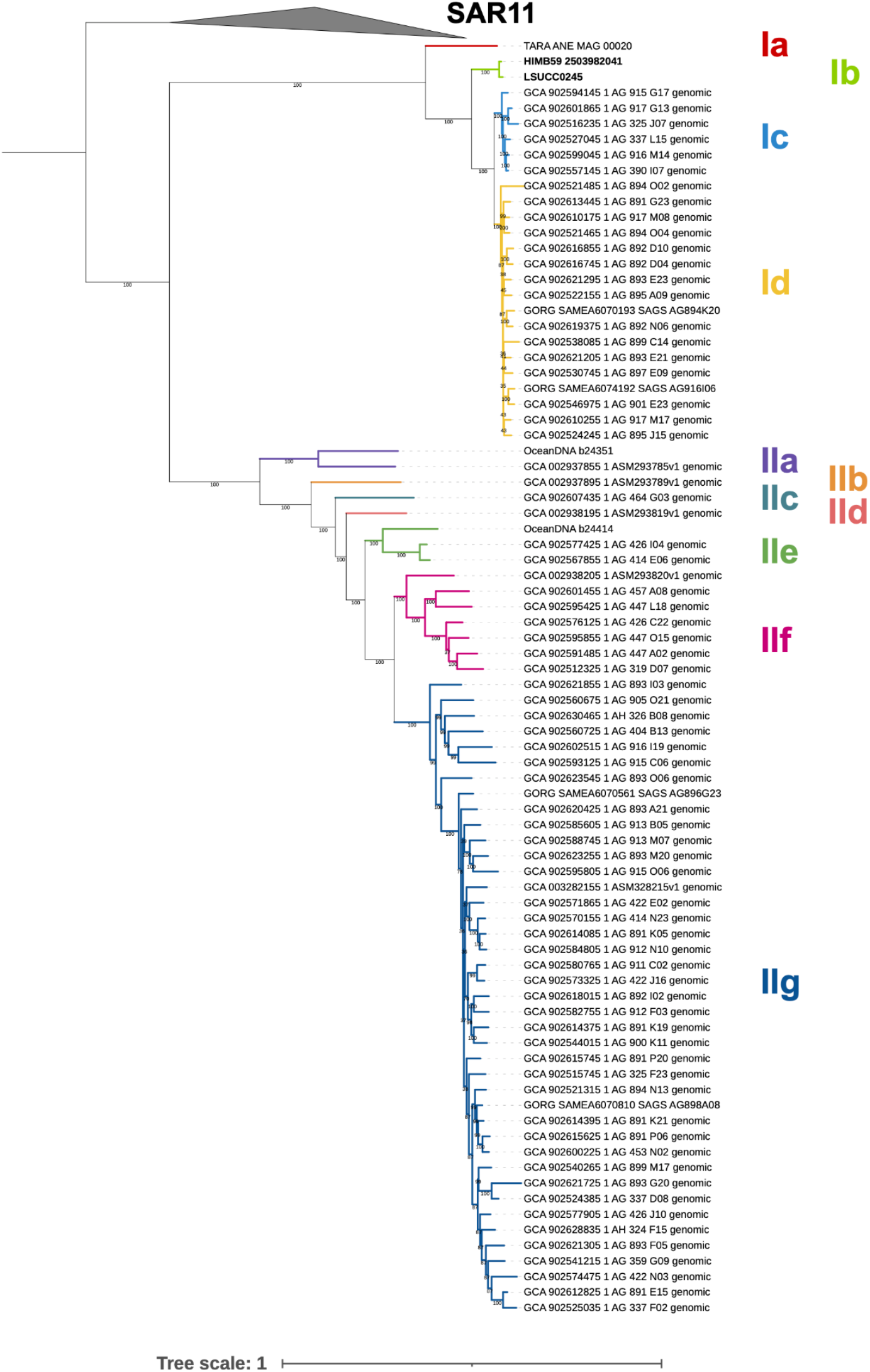
Phylogeny of AEGEAN-169 showing subclade designations. Values on the branches indicate ultrafast bootstrap support (n=1000), and subclade branches are colored to help provide contrast. Tree scale indicates changes per position according to the scale bar. SAR11 genomes were used as the outgroup.

### Genome Metrics

Estimated and actual genome sizes for AEGEAN-169 ranged from 1.26 Mbp to 1.84 Mbp with a mean of 1.55 Mbp (**Fig. 2**). The AEGEAN-169 genomes were larger than SAR11 (t-test, p<<0.01, R v4.2.1 (76)), which have genomes ranging 0.88 Mbp to 1.69 Mbp, with a mean of 1.22 Mbp). GC content for AEGEAN-169 ranged from 27.0% to 32.5% with a mean of 29.5%. These values were similar to SAR11 (t-test, p=0.09), whose GC content ranged from 27.6% to 35.9% with a mean of 29.3%. AEGEAN-169 coding densities ranged from 93.6% to 96.8% with a mean of 96.2%. SAR11 coding densities ranged from 92.0% to 97.1% with a mean of 96.4%. Thus, AEGEAN-169 had similar levels of genome streamlining to SAR11 even though the genomes were slightly larger.

**Figure 2.**
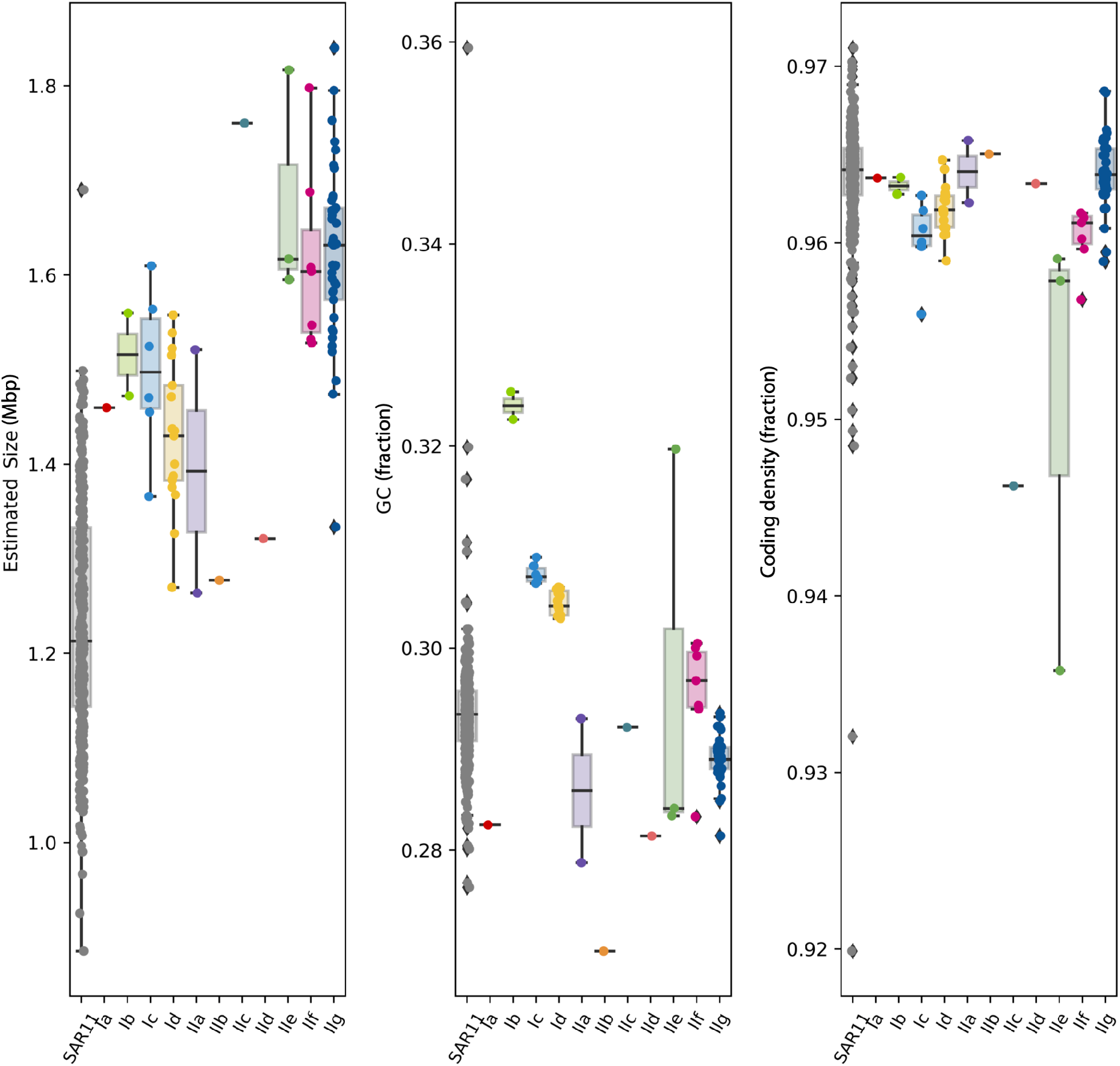
Boxplots illustrating the bulk genome characteristics of AEGEAN-169 subclades compared to SAR11. Subclades are colored according to the tree in Figure 1. Boxes describe the interquartile range (IQR) with the median indicated as a bar. Whiskers indicate 1.5x IQR and outlier points are plotted beyond the whiskers. The underlying datapoints for each boxplot is also plotted on top.

### Ecology

AEGEAN-169 is predominantly a surface water organism within the euphotic zone, with subgroup IIg dominating metagenomic recruitment in most marine locations, followed by subgroup Id (**Fig. S3a**). Subgroup IIc appeared to be a deep water bathytype, recruiting reads almost exclusively below 125m, with highest recruitment below the euphotic zone. Subgroup IIe was also more abundant in deeper waters, although it could be found at the surface (**Fig. S3a**). These patterns were consistent with distributions by temperature, where the surface subclades dominated in warmer temperatures, and the deeper subclades recruited most reads in colder water (**Fig. S3b**). We classified salinity according to the Venice system (< 0.5 fresh, 0.5-4.9 oligohaline, 5-17.9 mesohaline, 18-29.9 polyhaline, 30-39.9 euhaline, > 40 hyperhaline) (ITO 1959 (77)), confirming subgroup IIg as a marine organism with recruitment almost exclusively in euhaline and hyperhaline (**Fig. S3c**). Subgroup Ib was most prominent in polyhaline samples and recruited the most reads from mesohaline samples, so this likely represents a brackish water clade. None of the genomes within any subgroup represented freshwater taxa.

We also examined spatio-temporal trends from the Hawaii Ocean Time series (HOT) using samples collected monthly during the years 2003-2016 and normalizing by month. These samples extended to 500m. These data indicated that AEGEAN-169 has two primary ecological niches at HOT; surface water subgroups that bloom in the late summer/early fall and subgroups that occur primarily at 100-200 m and appear to have a fall bloom period (**Fig. 3)**. Subgroups Ic, Id, and IIg were the primary surface water groups, with Id and IIg being the most abundant at HOT, consistent with our global recruitment data (**Fig. S3**). Subgroups IIa, IId, and IIf were the dominant ecotypes in the 100-200 m range, suggesting they are associated with the deep chlorophyll maxima. Subgroups IIc and IIe were the only clades detected at 500 m, consistent with these organisms being deep water bathytypes.

**Figure 3.**
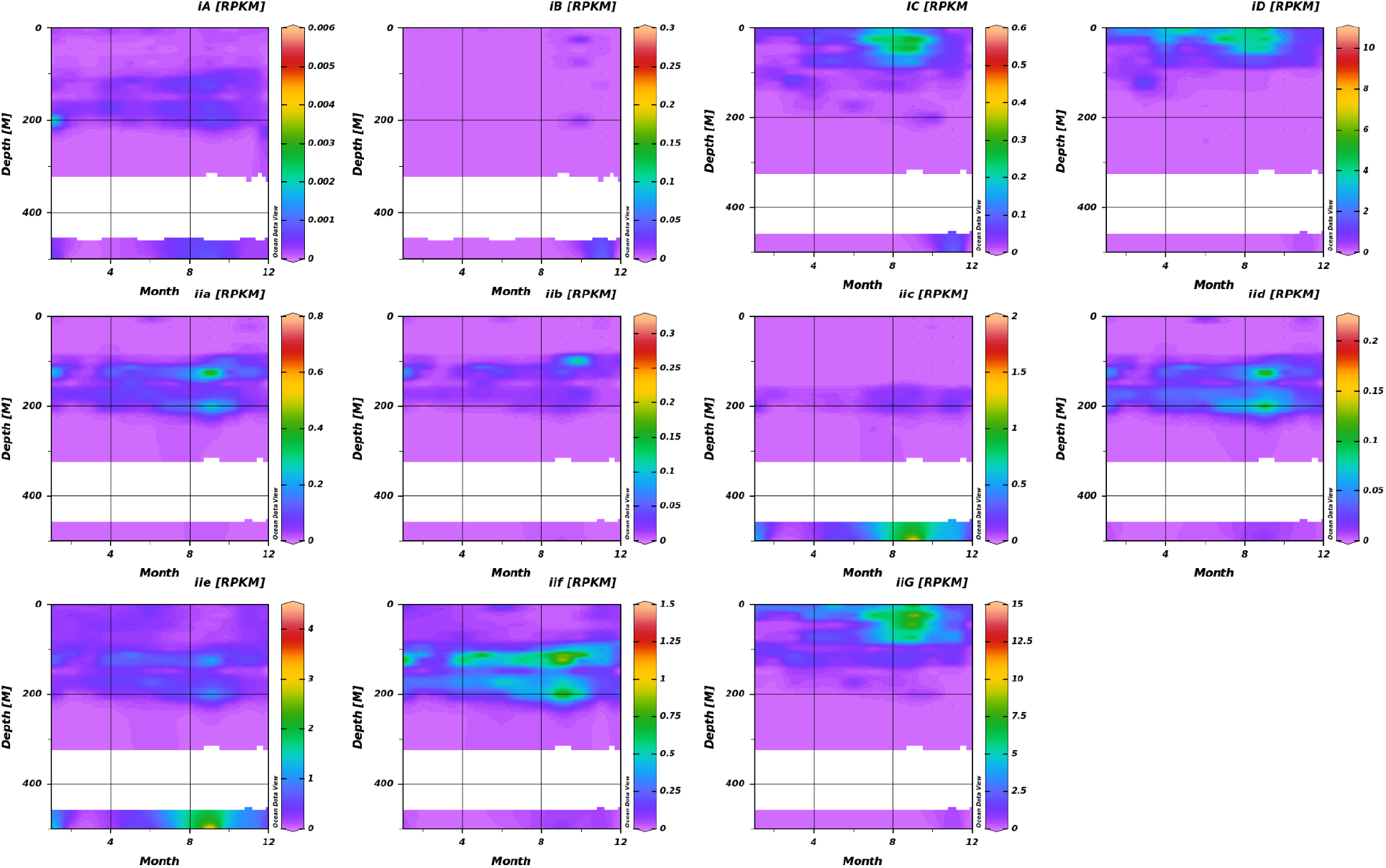
AEGEAN-169 subclade distribution at Station ALOHA using HOT data spanning from 2003-2016. Each subclade is plotted with a separate scale. Months correspond from 1 (January) to 12 (December). RPKM -Reads per kilobase (of genome) per megabase (of metagenome).

### Metabolic variation

What is currently known about the metabolism of AEGEAN-169 comes primarily from the HIMB59 genome (3). We have extended these observations to a larger diversity of genomes spanning the two subclades of AEGEAN-169. In general, these organisms are predicted to be obligate aerobes with chemoorganoheterotrophic metabolism. They have genes for central carbon metabolism by way of glycolysis, the pentose phosphate pathway, and the citric acid cycle, similar to SAR11. However, AEGEAN-169 metabolic capacity differs in several important ways, notably through sugar metabolism and trace metal and vitamin transport. AEGEAN-169 genomes had a fructose ABC transporter, predominantly in subclade I, subclade II members had a predicted trehalose/maltose ABC transporter, and both subclades included representatives with a galactose/raffinose/stachyose/mellibose ABC transporter-systems that are not found in SAR11 (**Fig. 4**). Although AEGEAN-169 genomes lacked an L-proline symporter found in SAR11, they shared the *potABCD* putrescine/spermidine transporter with SAR11, and had an additional *potFGBI* putrescine transporter and *algEFG* alpha-glucoside transporter not found in SAR11 (**Fig. 4**). Moreover, both AEGEAN-169 subclades had greater transport potential for trace metals and vitamins. Both subclades had heme and tungsten transporters not found in SAR11, and as well as the potential for thiamin transport that is absent in SAR11 (**Fig. 4**).

**Figure 4.**
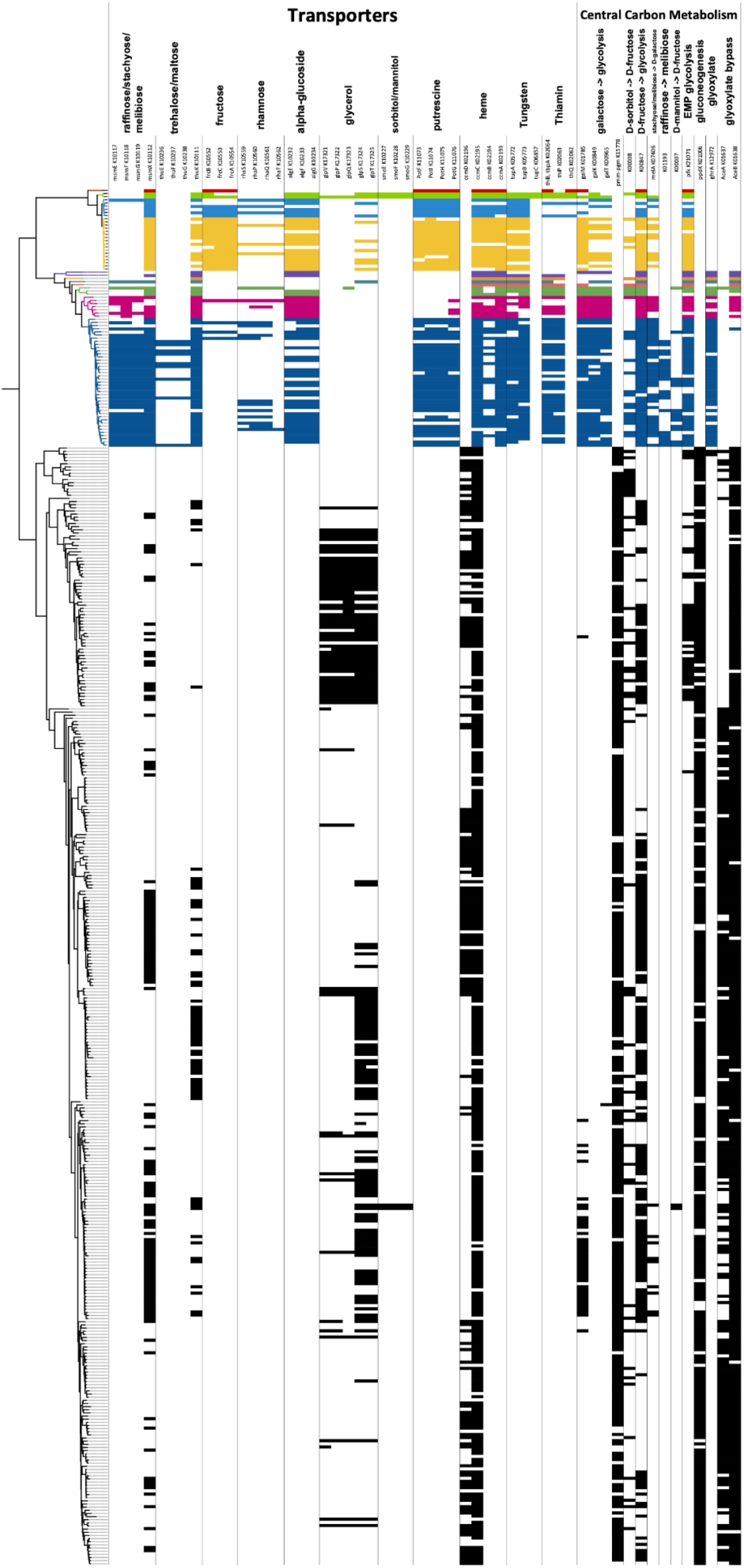
Key metabolic variation between AEGEAN-169 and SAR11. The phylogenomic tree on the left corresponds to the colors in Figure 1, with SAR11 indicated in only black branches below AEGEAN-169. Gene names correspond to components found for these systems.

AEGEAN-169 glycolytic inputs and central carbon metabolism also had key differences from those in SAR11. As reported previously for HIMB59 (3), AEGEAN-169 has the phosphofructokinase (pfk) for Embden–Meyerhof–Parnas glycolysis (**Fig. 4**). While this gene is found in some SAR11, including LD12 (78), it is missing from the dominant SAR11 subclade Ia organisms (**Fig. 4**). Consistent with the transporters for sugars, sugar metabolism was expanded. AEGEAN-169 members had predicted genes for the conversion of many sugars into galactose and/or fructose, as well as the *galKMT* pathway for galactose metabolism (**Fig. 4**). AEGEAN-169 also differed from SAR11 through the absence of *ppdK*, which converts phosphoenolpyruvate to pyruvate for gluconeogenesis (**Fig. 4**). While some subclade II members had *aceB* (malate synthase), we only found two examples of *aceA* (isocitrate lyase) in AEGEAN-169, and thus they appear to mostly lack the traditional glyoxylate shunt that is a hallmark of SAR11 (3, 78). However, most members of AEGEAN-169 subclade II had a predicted *ghrA* (glyoxylate/hydroxypyruvate reductase) (**Fig. 4**), which can convert glycolate to glyoxylate. Only two LD12 genomes had this gene within SAR11. AEGEAN-169 organisms with both *ghrA* and *aceB* should have the ability to bring glycolate into the TCA cycle, allowing them to take advantage of that widely abundant phytoplankton-produced compound (79, 80).

### Proteorhodopsin

We identified multiple gene clusters annotated as potential rhodopsin homologs within the pangenome, and numerous AEGEAN-169 genomes, including LSUCC0245, contain multiple copies of predicted proteorhodopsins (**Fig. S4**). Phylogenetic evaluation of these gene copies indicated multiple clusters with differential spectral tuning and a separate group of possible rhodopsin homologs that currently do not have functional prediction. These observations corroborate a recent investigation of proteorhodopsin paralogs in SAR11 and HIMB59-clade organisms (81).

### Physiology

We measured the growth rates of LSUCC0245 across multiple salinities and temperatures. This strain was a marine-adapted mesophile, growing optimally at 24°C, and slowly at 30°C, but not at 12 or 35°C (**Fig. 5a**). It grew optimally at a seawater salinity of 34 and in salinities as low as 11.6. Its maximum growth rate was 0.02 ± 0.007 divisions hr^-1^ at 24°C (**Fig. 5b**). LSUCC0245 had very low growth yields in our media (< 10^6^ cells ml^-1^) (**Fig. S5**). Given the complex mixture of low concentration carbon sources in the medium, it appears likely that LSUCC0245 was only using a small subset of the available substrates. We also note that several of the sugar, sugar alcohol, and polyamine compounds that we predict as usable by LSUCC0245 (e.g., sorbitol, mannitol, fructose, galactose, putrescine-**Fig. 4**) were not available in the JW2 medium (32). Thus, more in-depth exploration of usable carbon sources is warranted.

**Figure 5a.**
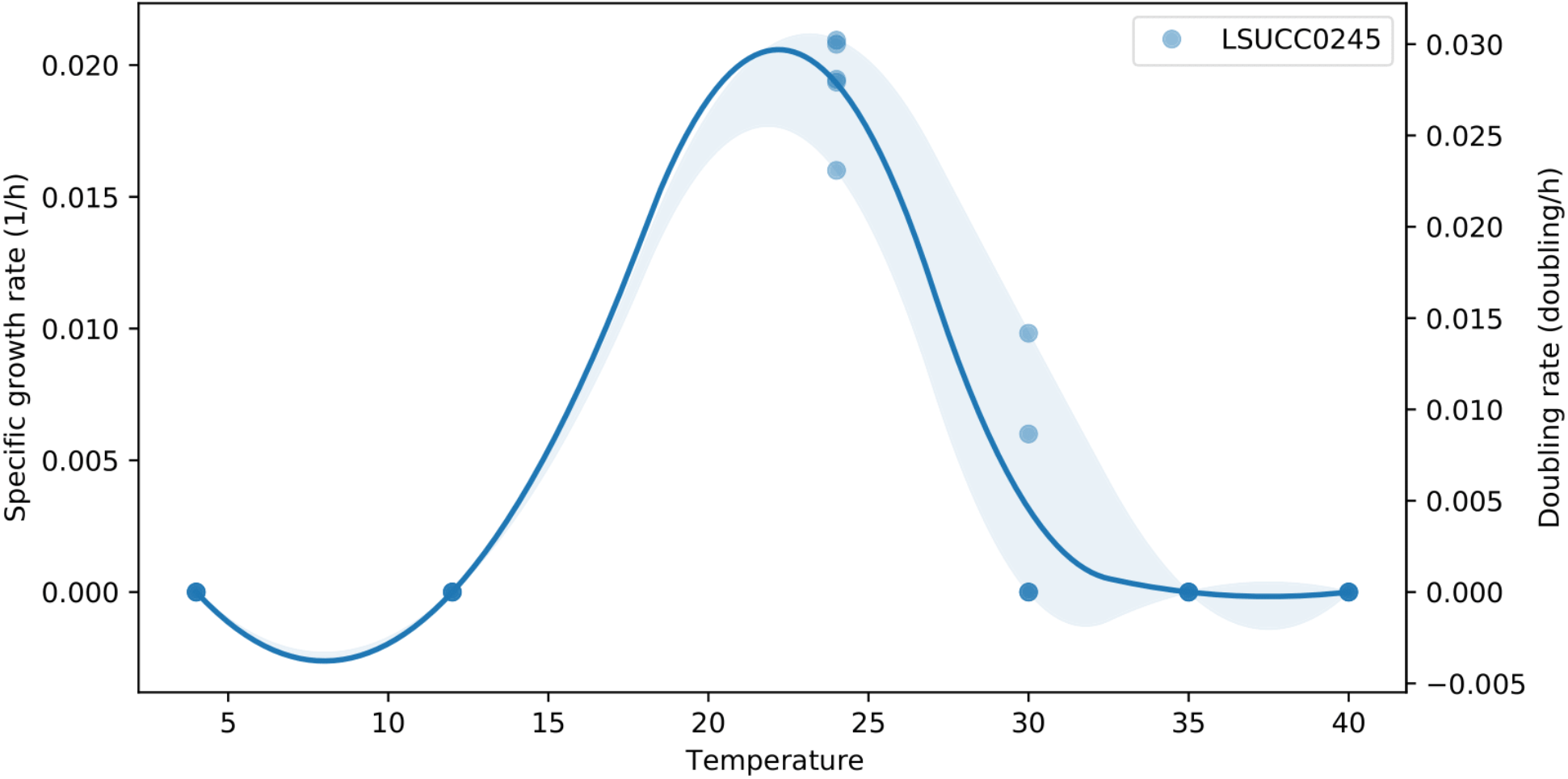
LSUCC0245 temperature dependent growth. Calculated using sparse-growth-curve (98). Specific growth rate and doubling rate are indicated with the dual y-axes. A best-fit line connects the points to predict rates in between measured values and shading indicates 95% confidence intervals.

**Figure 5b.**
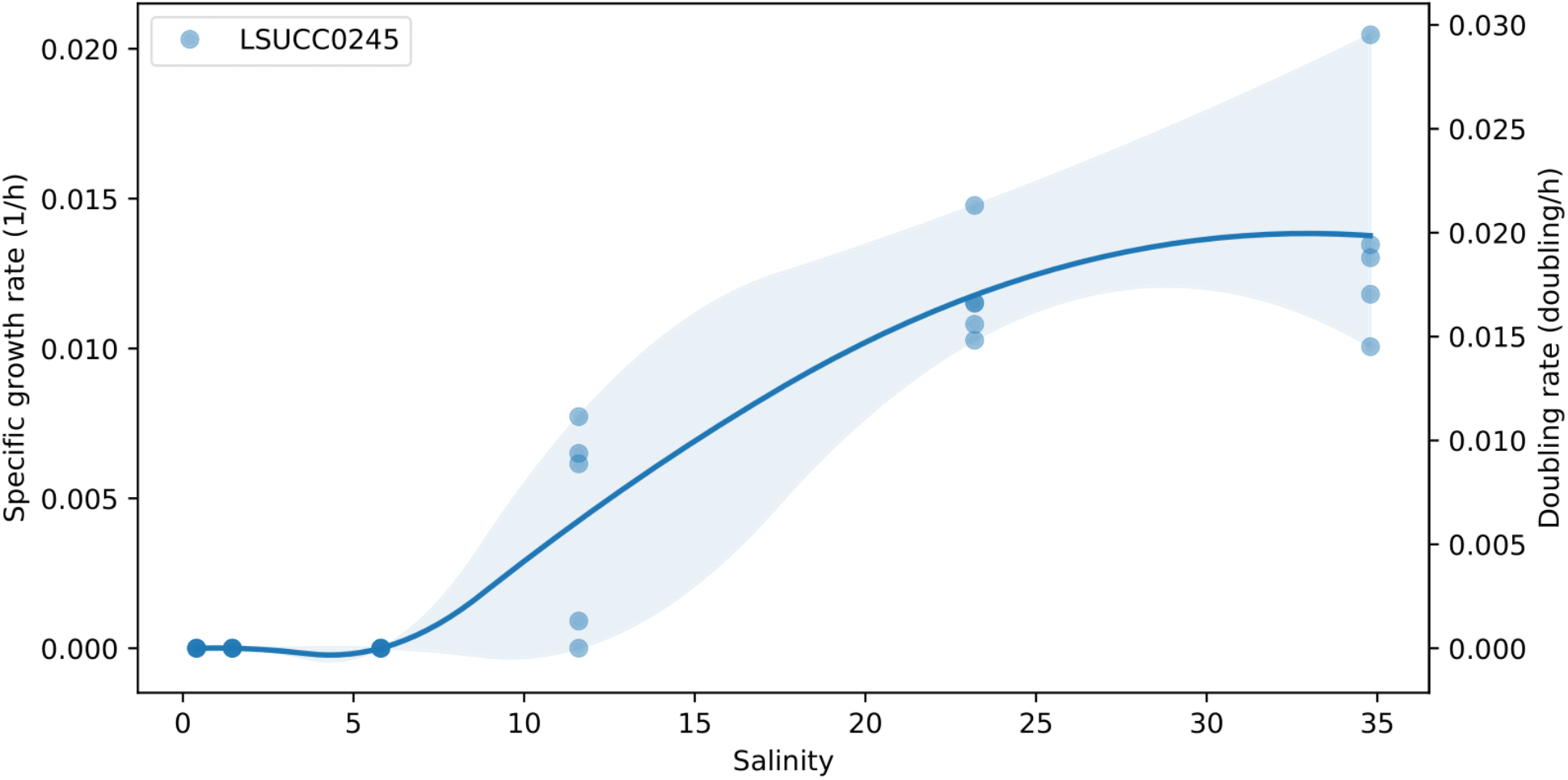
LSUCC0245 salinity dependent growth. Calculated using sparse-growth-curve (98). Specific growth rate and doubling rate are indicated with the dual y-axes. A best-fit line connects the points to predict rates in between measured values and shading indicates 95% confidence intervals.

## DISCUSSION

This study aimed to define AEGEAN-169 through the lens of taxonomy, ecology, and metabolism, with the goal of understanding how similar or distinct these organisms are from SAR11. Our results provide the first detailed examination of AEGEAN-169 genomics and genome-based ecology. The overall picture is one of a group that shares a very similar ecological regime as SAR11 – the majority of AEGEAN-169 members are most abundant in surface marine waters with seasonality that overlaps with SAR11. AEGEAN-169 and SAR11 were similar in relation to central carbon metabolism with a few key differences in capability. However, there were important metabolic differences between these groups, particularly the utilization of additional sugars, trace metals, and vitamins by AEGEAN-169, that may help distinguish their niche in terms of interactions with dissolved organic matter.

Although previous phylogenetic studies have considered SAR11 and AEGEAN-169 sister clades (21, 27, 82) this relationship likely results from compositional artifacts in the underlying sequence data (12, 13, 15). We reemphasize that our goal with phylogenetics and phylogenomics in this study was only to establish the subclade relationships within AEGEAN-169. Our work demonstrates, using both 16S rRNA genes and whole genome data, that AEGEAN-169 is a heterotopic synonym with SAR11 subclade V/HIMB59 (**Fig. S1**), thus condensing these disparate taxonomic designations. We propose using AEGEAN-169 as the primary moniker as we have done herein until a formal taxonomic designation is established. The major AEGEAN-169 subclade I and II delineation corresponds to the Va and Vb designations made on the basis of previous 16S rRNA gene phylogeny, respectively (7). While early work suggested a closer taxonomic relationship between HIMB59 and SAR11 through the use of synteny and genome organization (3), these observations were based on a singular genomic representative from AEGEAN-169 subclade I (HIMB59) and did not define the depth of the genus, as is now possible with current datasets (21, 28, 29). Thus, future examination of the phylogenetic relationships between AEGEAN-169, SAR11, and other Alphaproteobacteria should benefit from the expanded taxon selection provided by these studies.

Members of AEGEAN-169 were primarily surface-water marine organisms, sharing similar ecological distributions with SAR11 (7, 83–85). AEGEAN-169 was most abundant at depths between 5.1-75m. These observations corroborate previous characterizations of AEGEAN-169 as a predominantly surface water group that is likely stimulated by blooms occurring in late summer and fall (16, 18, 19, 86). AEGEAN-169 subgroups IIc and IIe were found predominantly in deeper waters (**Fig. S3a**) supporting bathytype designations, similarly to SAR11 subclade Ic (10). Notably, subclade II was the primary group with recruitment observed at 200 m or below, which is consistent with its distribution based on 16S rRNA gene data at BATS where subclade I (Va) was a surface water group, whereas subclade II (Vb) was found in both surface and 200 m waters (7). With respect to salinity, AEGEAN-169 were almost all marine-adapted, although subgroup Ib was most abundant in polyhaline conditions, suggesting that members of this subgroup inhabit a brackish niche (**Fig. S3c**) (12). The specific subgroup salinity preference resembles that described for SAR11 subclade IIIa (33). Overall, the high amount of overlap between the habitats of SAR11 and AEGEAN-169 likely explains the metabolic variation we observed between the two groups.

AEGEAN-169 lacked *ppdK* which converts phosphoenolpyruvate to pyruvate as well as the converse reaction (**Fig. 4**). This suggests that gluconeogenic activity is limited, which differs from the predicted complete gluconeogenesis pathway in SAR11 (3). Novel sugar intake was exhibited in AEGEAN-169 by means of multi-alpha-glucoside, fructose, rhamnose, trehalose/maltose, and raffinose/stachyose/mellibiose ABC transporters (**Fig. 4**). Expanded sugar metabolism was a feature first reported for HIMB59 based on the single genome at the time (3) and we demonstrate that this trait is conserved across AEGEAN-169 genomes. Many of the aforementioned sugars were predicted to be metabolized to galactose and through the *galKMT* genes (missing in SAR11) to alpha-D-glucose-1P (**Fig. 4**). However, AEGEAN-169 was missing the phosphoglucomutase found in SAR11, and we found no other means to convert alpha-D-glucose-1P to alpha-D-glucose-6P. Thus, how these sugars enter glycolysis is currently unclear. Nevertheless, given the greater emphasis on sugar transport and metabolism, but the lack of *ppdK*, perhaps AEGEAN-169 organisms rely on external sources of sugar instead of gluconeogenesis.

Another difference was that most AEGEAN-169 members had a predicted putrescine ABC transporter (*potFGHI*) not found in SAR11 (**Fig. 4**). SAR11 and some AEGEAN-169 members have homologs of the *potABCD* spermidine/putrescine ABC transporter (Supplemental Information -https://doi.org/10.6084/m9.figshare.22027763), and SAR11 responds disproportionately to addition of both of these polyamines in natural communities (87, 88). The *potABCD* genes transport five different polyamines in SAR11, where these compounds can meet cellular nitrogen requirements (89), and is spermidine-preferential in *Escherichia coli* (90). The additional *potFGHI* genes in AEGEAN-169 suggest increased use of putrescine compared to SAR11, as this transporter is considered putrescine-specific (90). Thus, SAR11 and AEGEAN-169 may have differential polyamine preferences in nature.

Trace metal and vitamin transport also distinguised AEGEAN-169 from SAR11. AEGEAN-169 uniquely had genes for an iron/zinc chelator, as well as heme and tungsten transport (**Fig. 4**). SAR11 members have quite limited trace metal transport capabilities (91). The potential of AEGEAN-169 to transport heme would provide them with an alternative source of iron, and the presence of the transporter corroborates recent findings that many abundant marine microorganisms are heme auxotrophs (92), including AEGEAN-169 members (designated HIMB59 by the authors). Most surprising was the presence of a tungsten transporter which traditionally has been observed in thermophilic archaea as well as *Sulfitobacteria dubius* and some *Clostridium* spp. and *Eubacterium* spp., although hyperthermophilic archaea appear to be the only group that requires tungsten (93–95). This suggests that AEGEAN-169 may utilize tungstoenzymes, such as a tungsten-containing version of formate dehydrogenase (96). Formate deydrogenases are conserved throughout SAR11 and AEGEAN-169 (Supplemental Information -https://doi.org/10.6084/m9.figshare.22027763), but the clades may use different cofactors. AEGEAN-169 also had the capacity to transport thiamin (vitamin B1), which may provide another means of niche differentiation since SAR11 relies on thiamin precursors instead of directly uptaking thiamin (97).

The increased potential of sugar, trace metal, and vitamin transport and metabolism are important traits differentiating AEGEAN-169 from SAR11, and likely mean that AEGEAN-169 has a more extensive metabolic niche than SAR11. This expanded metabolic repertoire correlates with the slightly larger genome sizes in AEGEAN-169 compared to SAR11, even though both strains have the hallmark coding density associated with genome streamlining. Nevertheless, SAR11 is the more successful group, with relative abundances that are usually much higher than that of AEGEAN-169 (e.g., (7)). In this context, it is notable that strain LSUCC0245 grew to much lower cell densities than SAR11 strains in the same medium, even though growth rates were similar (33, 78). Since our defined media have numerous carbon compounds at similar concentrations, these yield differences either mean that SAR11 and AEGEAN-169 use a different set of compounds available in the medium, or there is something inherently different about growth physiology with AEGEAN-169. Future studies should incorporate cultivation assessments to investigate the metabolic differences we have identified, as well as the differences in physiology. Additional isolates will also help improve our overall understanding of the diversity of functions in the group and shed more light on the evolutionary pressures that have led to the similarities that AEGEAN-169 and SAR11 share.

## Data Availability

Raw reads for the LSUCC0245 genome were deposited at NCBI BioProject number <u>PRJNA931292</u>, and the genome is publicly available on IMG (https://img.jgi.doe.gov/) under Genome ID: 2756170191. Supporting datasets, scripts, and files are available at https://doi.org/10.6084/m9.figshare.22027763.

## Acknowledgements

We thank Oded Beja for insightful comments and reference sequences for rhodopsins. The authors acknowledge the Center for Advanced Research Computing (CARC) at the University of Southern California for providing computing resources that have contributed to the research results reported within this publication. URL: https://carc.usc.edu. Portions of this research were conducted with high performance computing resources provided by Louisiana State University (http://www.hpc.lsu.edu). This work was supported by a Simons Early Career Investigator in Marine Microbial Ecology and Evolution Award, and NSF Biological Oceanography Program OCE-1945279 and Emerging Frontiers Program EF-2125191 grants to J.C.T.

---

**Figure S1.**
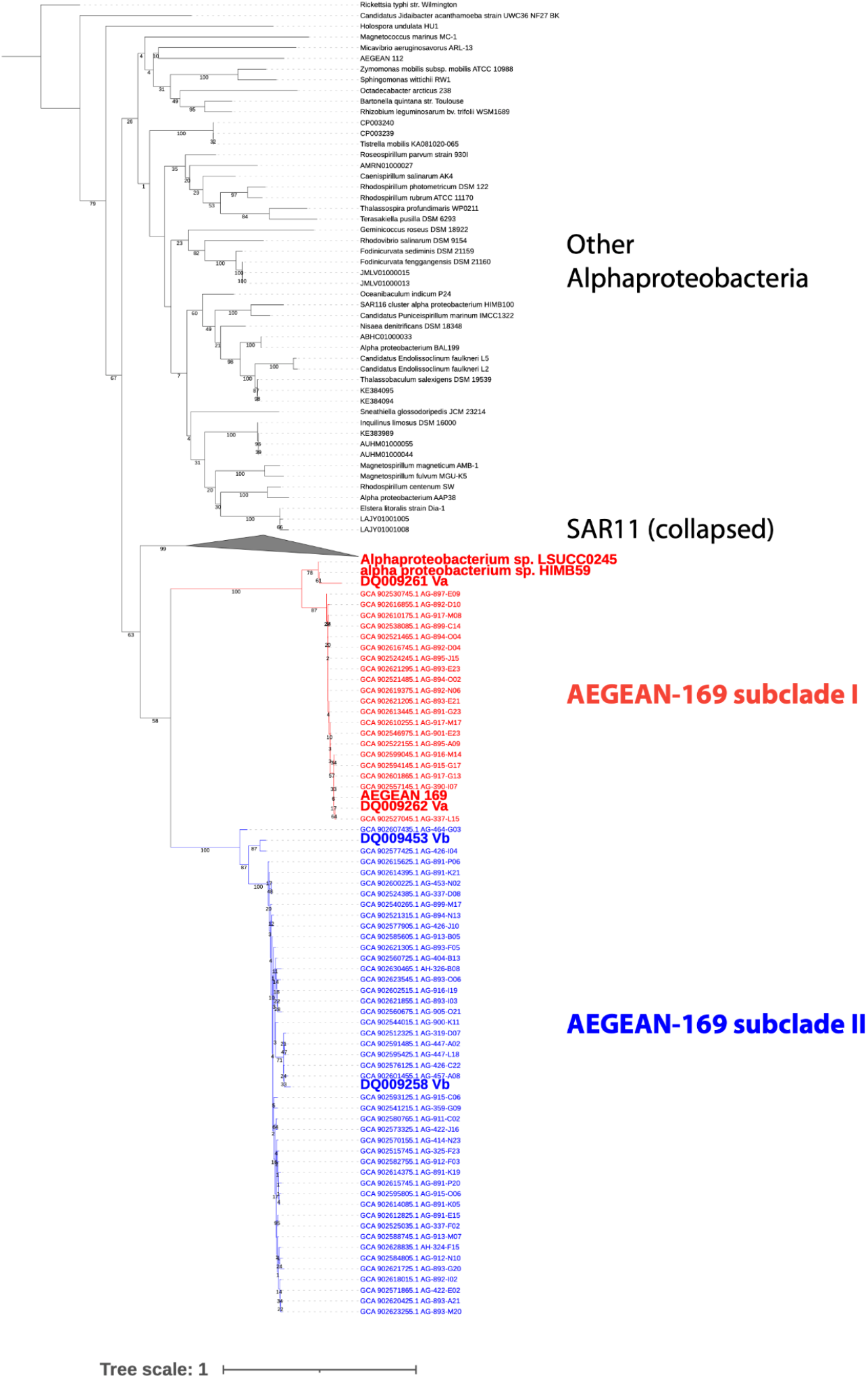
16S rRNA gene tree phylogeny of AEGEAN-169, SAR11, and other Alphaproteobacteria. Genomes and clone library markers of interest are bolded within the AEGEAN-169 subclades for emphasis. All sequences for SAR11 are collapsed. The original AEGEAN-169 16S rRNA gene marker groups with the SAR11 subclade Va markers and the two cultured isolates, HIMB59 and LSUCC0245 in subclade I. The SAR11 subclade Vb markers group with AEGEAN-169 subclade II. Values at the nodes indicate traditional bootstraps (n=100) and the Tree scale indicates changes per position according to the bar. The tree was rooted on *Rickettsia typhi*.

**Figure S2.**
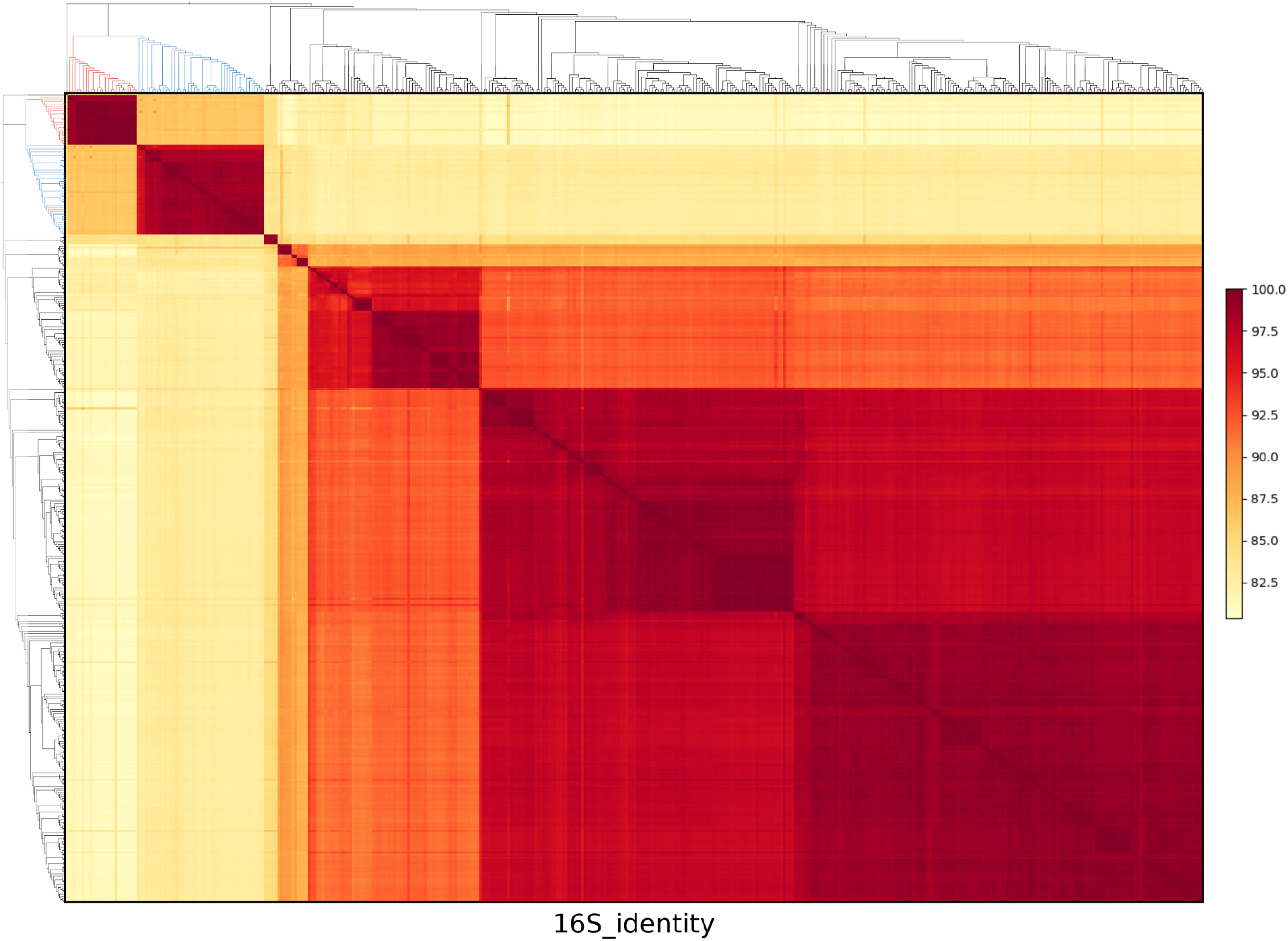
16S rRNA gene identity of AEGEAN-169 and SAR11. Subclade colors of the dendrogram correspond to those in Fig. S1, excluding other Alphaproteobacteria. Percent identity is denoted according to the scale bar on the right.

**Figure S3a.**
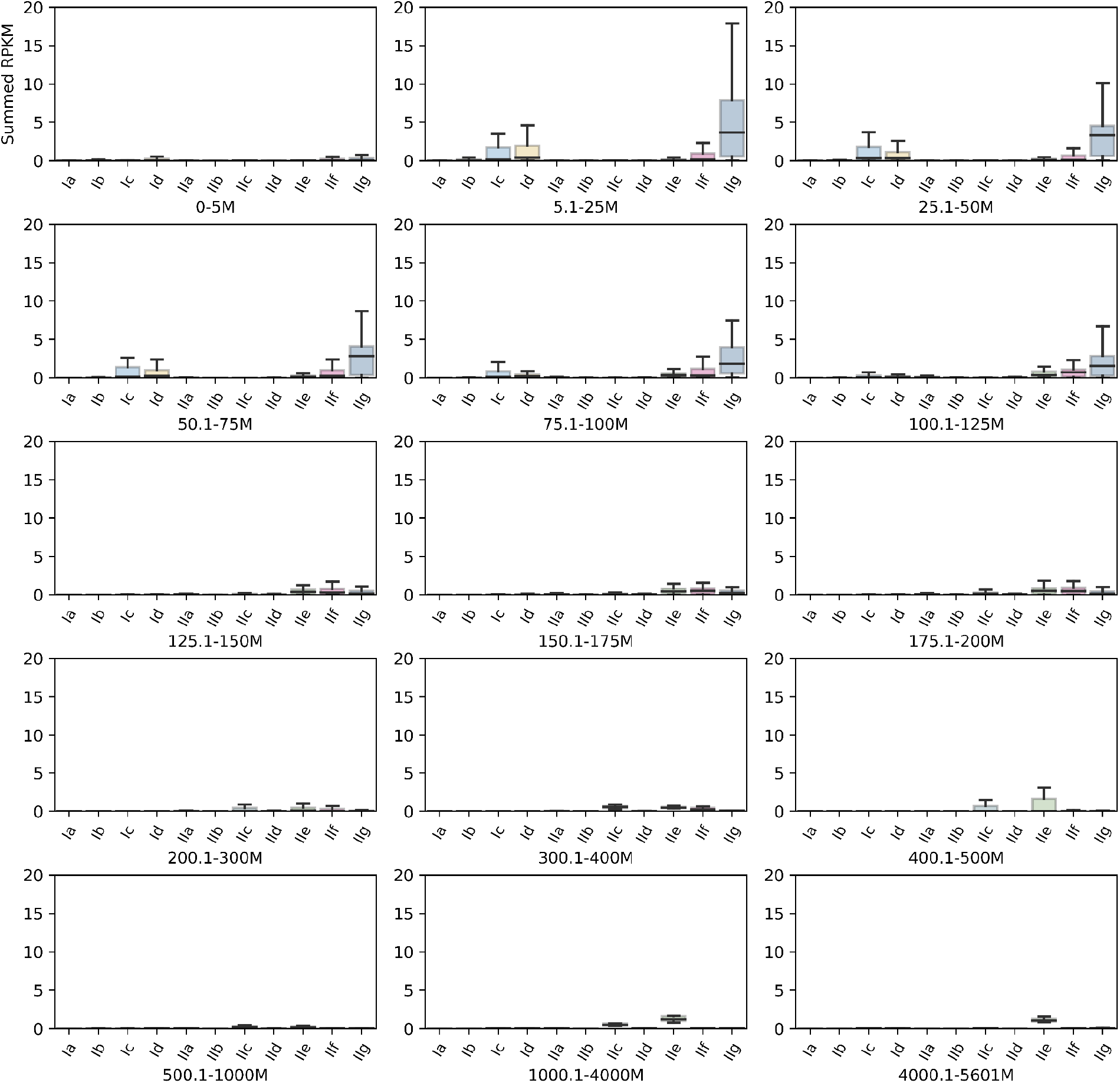
AEGEAN-169 subclade distribution by depth. Subclades are plotted according to the sum of the individual genome RPKMs comprising that subclade. RPKM - Reads per kilobase of genome sequence per megabase of metagenomic sequence.

**Figure S3b.**
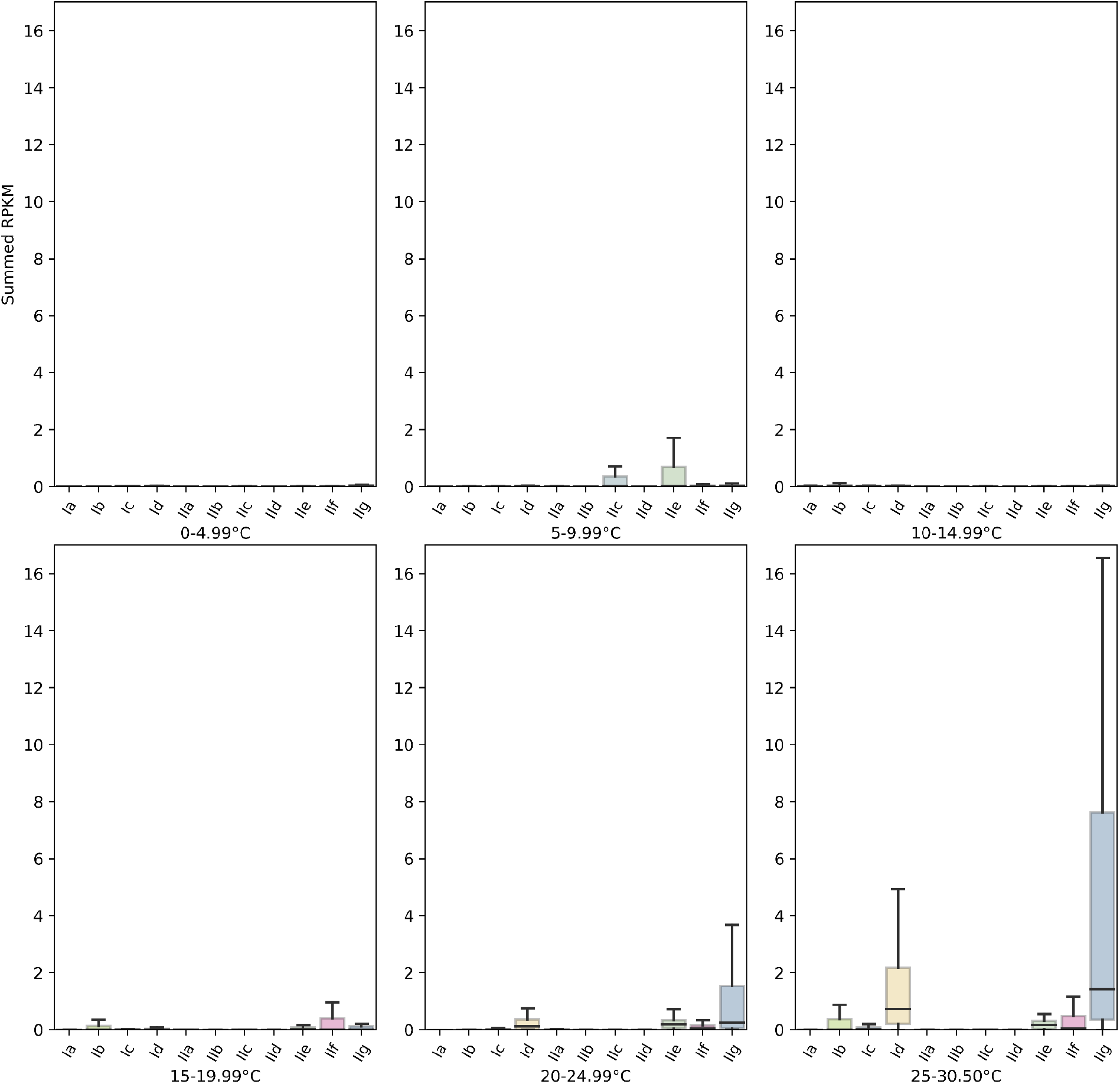
AEGEAN-169 subclade distribution by temperature. Subclades are plotted according to the sum of the individual genome RPKMs comprising that subclade. RPKM -Reads per kilobase of genome sequence per megabase of metagenomic sequence.

**Figure S3c.**
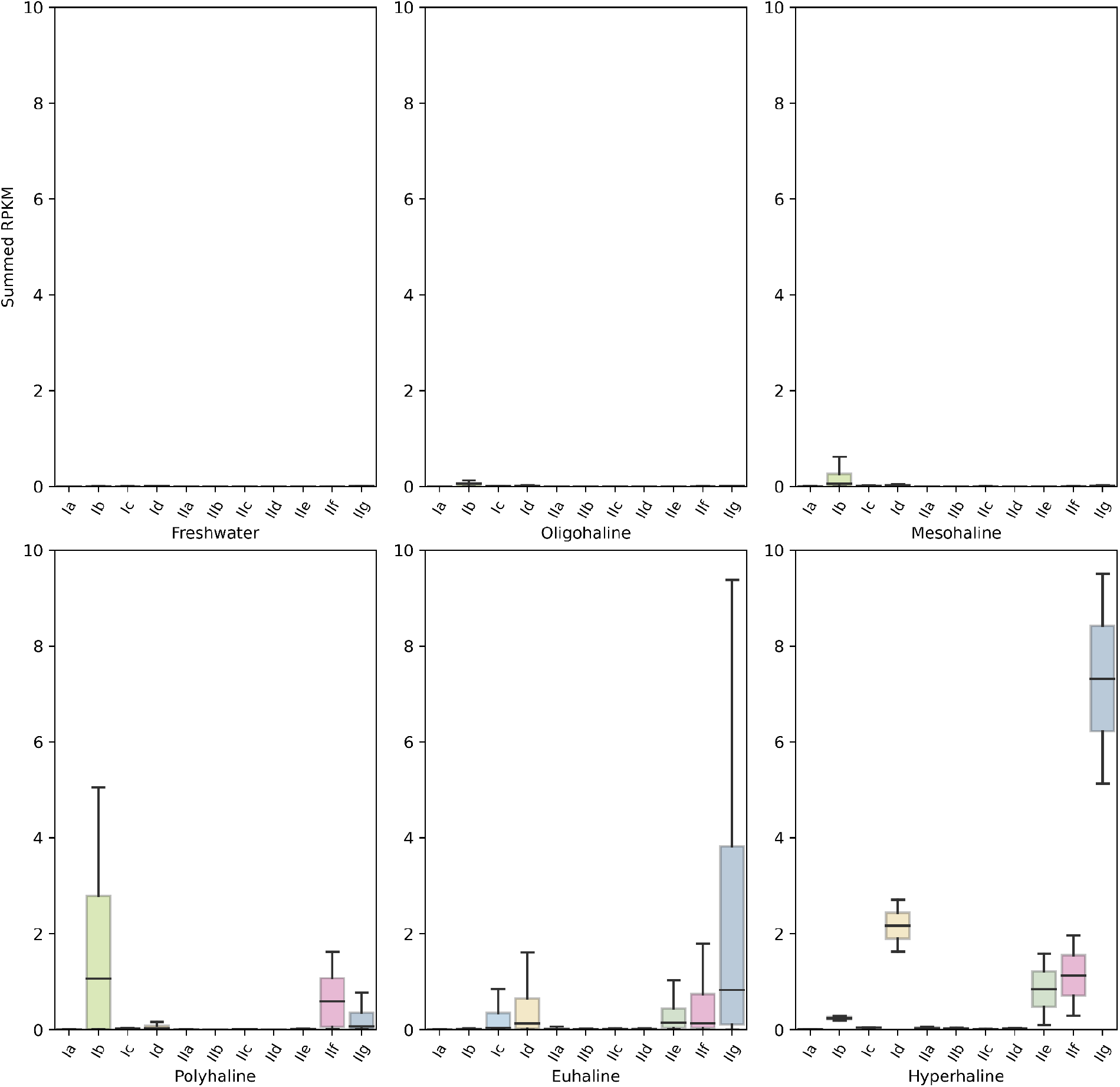
AEGEAN-169 subclade distribution by salinity. Subclades are plotted according to the sum of the individual genome RPKMs comprising that subclade. RPKM -Reads per kilobase of genome sequence per megabase of metagenomic sequence. Salinity categories are as follows: < 0.5 fresh, 0.5-4.9 oligohaline, 5-17.9 mesohaline, 18-29.9 polyhaline, 30-39.9 euhaline, > 40 hyperhaline

**Figure S4.**
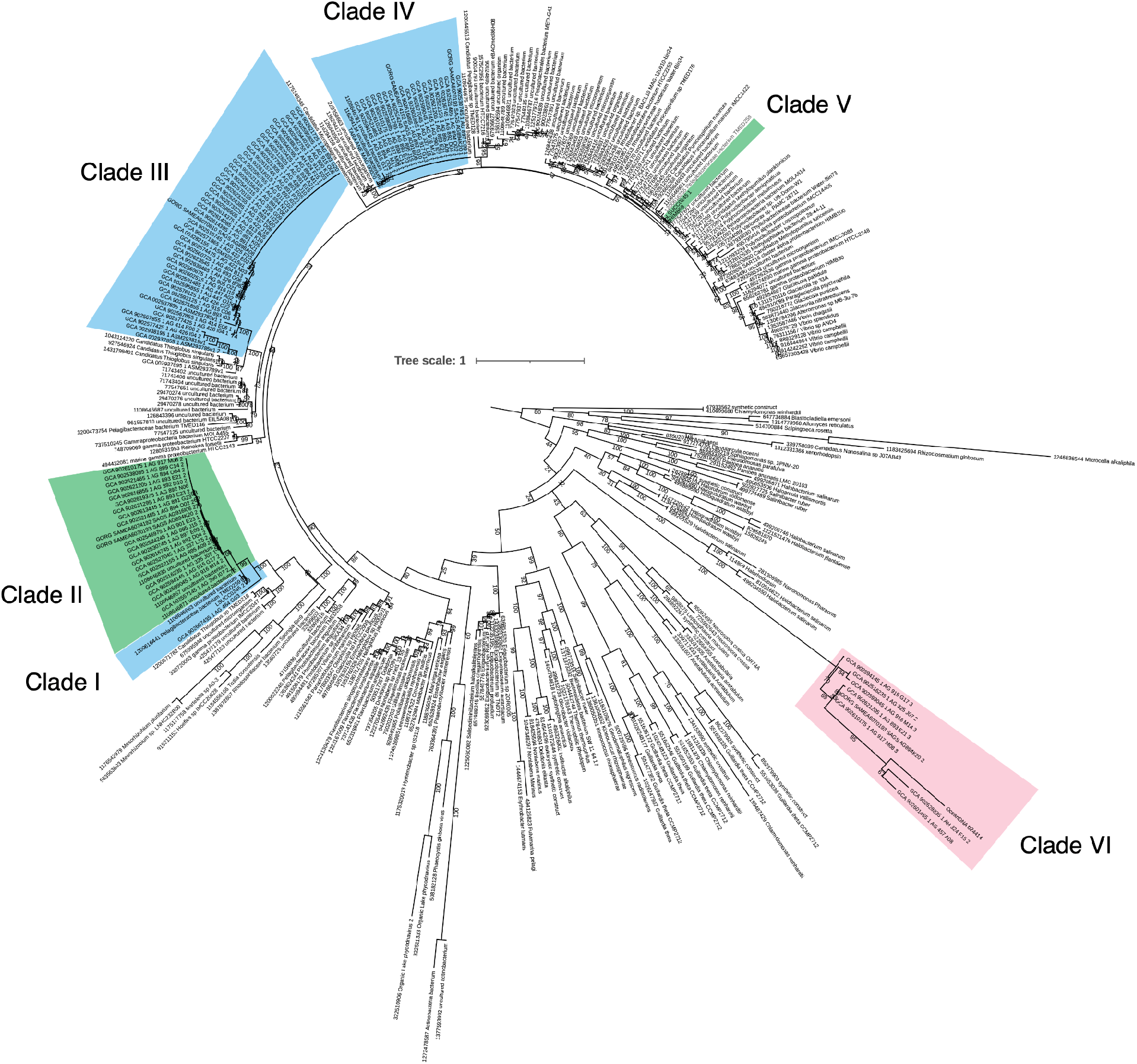
Proteorhodopsin phylogeny. Ultrafast bootstrap values (n=1000) are indicated on the branches and clades with AEGEAN-169 members are highlighted. Blue and green highlighting corresponds to spectral tuning of those groups, which are arbitrarily given Clade I-V names, whereas the red highlight for Clade VI indicates an undetermined function. Tree scale indicates changes per position according to the scale bar.

**Figure S5.**
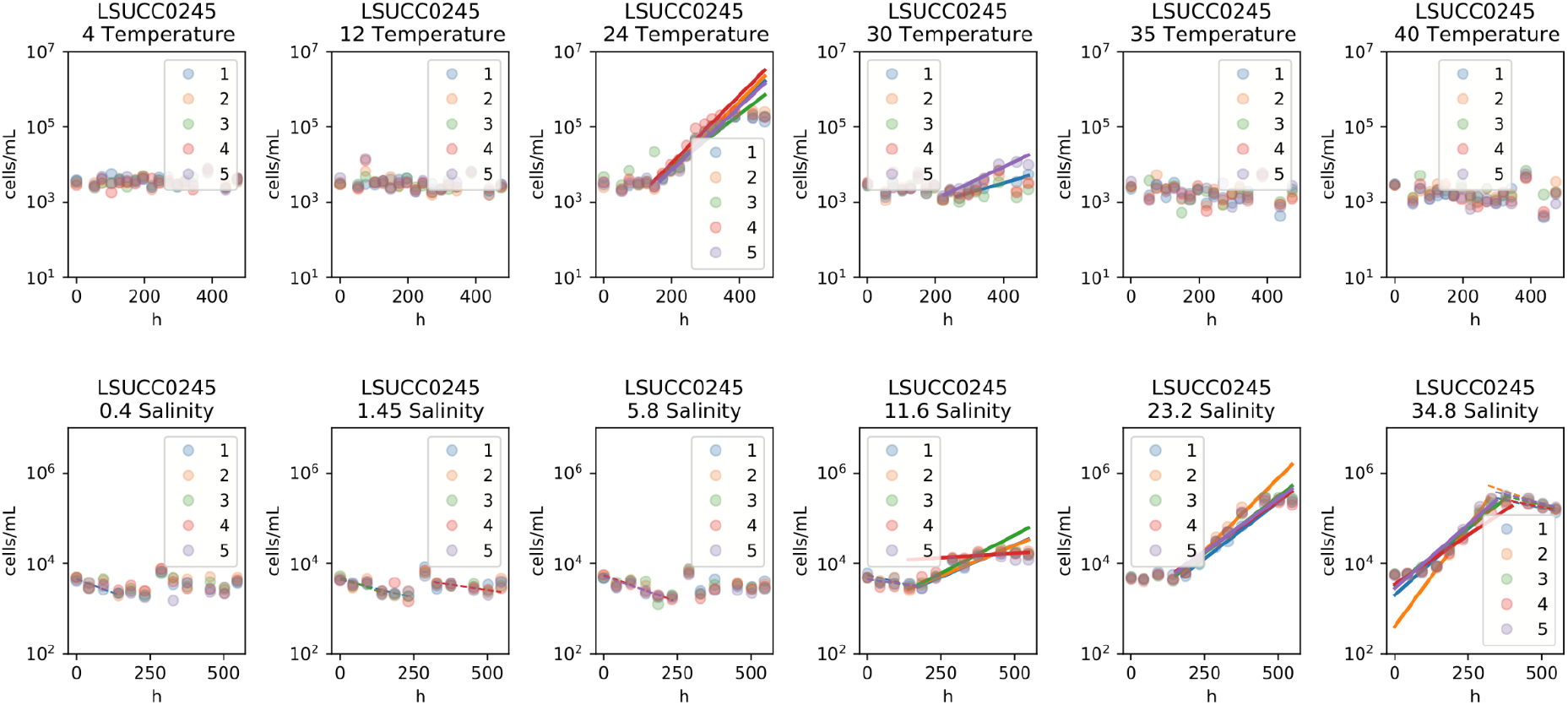
Growth data for LSUCC0245 temperature and salinity experiments. These data underlie the computed rates in Fig. 5.

## REFERENCES

1. Giovannoni SJ. 2017. SAR11 Bacteria: The Most Abundant Plankton in the Oceans. Ann Rev Mar Sci 9:231–255.

2. Giovannoni SJ, Tripp HJ, Givan S, Podar M, Vergin KL, Baptista D, Bibbs L, Eads J, Richardson TH, Noordewier M, Rappé MS, Short JM, Carrington JC, Mathur EJ. 2005. Genome streamlining in a cosmopolitan oceanic bacterium. Science 309:1242–1245.

3. Grote J, Thrash JC, Huggett MJ, Landry ZC, Carini P, Giovannoni SJ, Rappé MS. 2012. Streamlining and core genome conservation among highly divergent members of the SAR11 clade. MBio 3.

4. Giovannoni SJ, Cameron Thrash J, Temperton B. 2014. Implications of streamlining theory for microbial ecology. ISME J 8:1553–1565.

5. Giovannoni SJ, Vergin KL. 2012. Seasonality in ocean microbial communities. Science 335:671–676.

6. Haro-Moreno JM, Rodriguez-Valera F, Rosselli R, Martinez-Hernandez F, Roda-Garcia JJ, Gomez ML, Fornas O, Martinez-Garcia M, López-Pérez M. 2020. Ecogenomics of the SAR11 clade. Environ Microbiol 22:1748–1763.

7. Vergin KL, Beszteri B, Monier A, Thrash JC, Temperton B, Treusch AH, Kilpert F, Worden AZ, Giovannoni SJ. 2013. High-resolution SAR11 ecotype dynamics at the Bermuda Atlantic Time-series Study site by phylogenetic placement of pyrosequences. ISME J 7:1322–1332.

8. Ruiz-Perez CA, Bertagnolli AD, Tsementzi D, Woyke T, Stewart FJ, Konstantinidis KT. 2021. Description of Candidatus Mesopelagibacter carboxydoxydans and Candidatus Anoxipelagibacter denitrificans: Nitrate-reducing SAR11 genera that dominate mesopelagic and anoxic marine zones. Syst Appl Microbiol 126185.

9. Tsementzi D, Wu J, Deutsch S, Nath S, Rodriguez-R LM, Burns AS, Ranjan P, Sarode N, Malmstrom RR, Padilla CC, Stone BK, Bristow LA, Larsen M, Glass JB, Thamdrup B, Woyke T, Konstantinidis KT, Stewart FJ. 2016. SAR11 bacteria linked to ocean anoxia and nitrogen loss. Nature 536:179–183.

10. Thrash JC, Cameron Thrash J, Temperton B, Swan BK, Landry ZC, Woyke T, DeLong EF, Stepanauskas R, Giovannoni SJ. 2014. Single-cell enabled comparative genomics of a deep ocean SAR11 bathytype. The ISME Journal https://doi.org/10.1038/ismej.2013.243.

11. Ferla MP, Thrash JC, Giovannoni SJ, Patrick WM. 2013. New rRNA gene-based phylogenies of the Alphaproteobacteria provide perspective on major groups, mitochondrial ancestry and phylogenetic instability. PLoS One 8:e83383.

12. Viklund J, Martijn J, Ettema TJG, Andersson SGE. 2013. Comparative and phylogenomic evidence that the alphaproteobacterium HIMB59 is not a member of the oceanic SAR11 clade. PLoS One 8:e78858.

13. Martijn J, Vosseberg J, Guy L, Offre P, Ettema TJG. 2018. Deep mitochondrial origin outside the sampled alphaproteobacteria. Nature 557:101–105.

14. Muñoz-Gómez SA, Hess S, Burger G, Lang BF, Susko E, Slamovits CH, Roger AJ. 2019. An updated phylogeny of the Alphaproteobacteria reveals that the Rickettsiales and Holosporales have independent origins. Elife 8:e42535.

15. Moeseneder MM, Arrieta JM, Herndl GJ. 2005. A comparison of DNA-and RNA-based clone libraries from the same marine bacterioplankton community. FEMS Microbiol Ecol 51:341–352.

16. Yang C, Li Y, Zhou B, Zhou Y, Zheng W, Tian Y, Van Nostrand JD, Wu L, He Z, Zhou J, Zheng T. 2015. Illumina sequencing-based analysis of free-living bacterial community dynamics during an Akashiwo sanguine bloom in Xiamen sea, China. Sci Rep 5:8476.

17. Cram JA, Xia LC, Needham DM, Sachdeva R, Sun F, Fuhrman JA. 2015. Cross-depth analysis of marine bacterial networks suggests downward propagation of temporal changes. ISME J 9:2573–2586.

18. Reintjes G, Tegetmeyer HE, Bürgisser M, Orlić S, Tews I, Zubkov M, Voß D, Zielinski O, Quast C, Glöckner FO, Amann R, Ferdelman TG, Fuchs BM. 2019. On-Site Analysis of Bacterial Communities of the Ultraoligotrophic South Pacific Gyre. Appl Environ Microbiol 85.

19. Allen R, Hoffmann LJ, Law CS, Summerfield TC. 2020. Subtle bacterioplankton community responses to elevated CO2 and warming in the oligotrophic South Pacific gyre. Environ Microbiol Rep 12:377–386.

20. Šantić D, Piwosz K, Matić F, Vrdoljak Tomaš A, Arapov J, Dean JL, Šolić M, Koblížek M, Kušpilić G, Šestanović S. 2021. Artificial neural network analysis of microbial diversity in the central and southern Adriatic Sea. Sci Rep 11:11186.

21. Pachiadaki MG, Brown JM, Brown J, Bezuidt O, Berube PM, Biller SJ, Poulton NJ, Burkart MD, La Clair JJ, Chisholm SW, Stepanauskas R. 2019. Charting the Complexity of the Marine Microbiome through Single-Cell Genomics. Cell 179:1623–1635.e11.

22. van Bleijswijk JDL, Whalen C, Duineveld GCA, Lavaleye MSS, Witte HJ, Mienis F. 2015. Microbial assemblages on a cold-water coral mound at the SE Rockall Bank (NE Atlantic): interactions with hydrography and topography. Biogeosciences 12:4483–4496.

23. Korlević M, Markovski M, Herndl GJ, Najdek M. 2022. Temporal variation in the prokaryotic community of a nearshore marine environment. Sci Rep 12:16859.

24. Steiner PA, Sintes E, Simó R, De Corte D, Pfannkuchen DM, Ivančić I, Najdek M, Herndl GJ. 2019. Seasonal dynamics of marine snow-associated and free-living demethylating bacterial communities in the coastal northern Adriatic Sea. Environ Microbiol Rep 11:699–707.

25. Tong F, Zhang P, Zhang X, Chen P. 2021. Impact of oyster culture on coral reef bacterioplankton community composition and function in Daya Bay, China. Aquac Environ Interact 13:489–503.

26. Li Y-Y, Chen X-H, Xie Z-X, Li D-X, Wu P-F, Kong L-F, Lin L, Kao S-J, Wang D-Z. 2018. Bacterial Diversity and Nitrogen Utilization Strategies in the Upper Layer of the Northwestern Pacific Ocean. Front Microbiol 9:797.

27. Acker M, Hogle SL, Berube PM, Hackl T, Coe A, Stepanauskas R, Chisholm SW, Repeta DJ. 2022. Phosphonate production by marine microbes: Exploring new sources and potential function. Proc Natl Acad Sci U S A 119:e2113386119.

28. Paoli L, Ruscheweyh H-J, Forneris CC, Hubrich F, Kautsar S, Bhushan A, Lotti A, Clayssen Q, Salazar G, Milanese A, Carlström CI, Papadopoulou C, Gehrig D, Karasikov M, Mustafa H, Larralde M, Carroll LM, Sánchez P, Zayed AA, Cronin DR, Acinas SG, Bork P, Bowler C, Delmont TO, Gasol JM, Gossert AD, Kahles A, Sullivan MB, Wincker P, Zeller G, Robinson SL, Piel J, Sunagawa S. 2022. Biosynthetic potential of the global ocean microbiome. Nature 607:111–118.

29. Nishimura Y, Yoshizawa S. 2022. The OceanDNA MAG catalog contains over 50,000 prokaryotic genomes originated from various marine environments. Sci Data 9:305.

30. Parks DH, Chuvochina M, Rinke C, Mussig AJ, Chaumeil P-A, Hugenholtz P. 2021. GTDB: an ongoing census of bacterial and archaeal diversity through a phylogenetically consistent, rank normalized and complete genome-based taxonomy. Nucleic Acids Res https://doi.org/10.1093/nar/gkab776.

31. Henson MW, Lanclos VC, Pitre DM, Weckhorst JL, Lucchesi AM, Cheng C, Temperton B, Thrash JC. 2020. Expanding the Diversity of Bacterioplankton Isolates and Modeling Isolation Efficacy with Large-Scale Dilution-to-Extinction Cultivation. Appl Environ Microbiol 86.

32. Henson MW, Pitre DM, Weckhorst JL, Lanclos VC, Webber AT, Thrash JC. 2016. Artificial Seawater Media Facilitate Cultivating Members of the Microbial Majority from the Gulf of Mexico. mSphere 1.

33. Lanclos VC, Rasmussen AN, Kojima CY, Cheng C, Henson MW, Faircloth BC, Francis CA, Thrash JC. 2023. Ecophysiology and genomics of the brackish water adapted SAR11 subclade IIIa. ISME J https://doi.org/10.1038/s41396-023-01376-2.

34. Bolger AM, Lohse M, Usadel B. 2014. Trimmomatic: a flexible trimmer for Illumina sequence data. Bioinformatics 30:2114–2120.

35. Bankevich A, Nurk S, Antipov D, Gurevich AA, Dvorkin M, Kulikov AS, Lesin VM, Nikolenko SI, Pham S, Prjibelski AD, Pyshkin AV, Sirotkin AV, Vyahhi N, Tesler G, Alekseyev MA, Pevzner PA. 2012. SPAdes: a new genome assembly algorithm and its applications to single-cell sequencing. J Comput Biol 19:455–477.

36. Walker BJ, Abeel T, Shea T, Priest M, Abouelliel A, Sakthikumar S, Cuomo CA, Zeng Q, Wortman J, Young SK, Earl AM. 2014. Pilon: an integrated tool for comprehensive microbial variant detection and genome assembly improvement. PLoS One 9:e112963.

37. Li H, Durbin R. 2009. Fast and accurate short read alignment with Burrows-Wheeler transform. Bioinformatics 25:1754–1760.

38. Chen I-MA, Chu K, Palaniappan K, Pillay M, Ratner A, Huang J, Huntemann M, Varghese N, White JR, Seshadri R, Smirnova T, Kirton E, Jungbluth SP, Woyke T, Eloe-Fadrosh EA, Ivanova NN, Kyrpides NC. 2019. IMG/M v.5.0: an integrated data management and comparative analysis system for microbial genomes and microbiomes. Nucleic Acids Research https://doi.org/10.1093/nar/gky901.

39. Markowitz VM, Ivanova NN, Szeto E, Palaniappan K, Chu K, Dalevi D, Chen I-MA, Grechkin Y, Dubchak I, Anderson I, Lykidis A, Mavromatis K, Hugenholtz P, Kyrpides NC. 2008. IMG/M: a data management and analysis system for metagenomes. Nucleic Acids Res 36:D534–8.

40. Jain C, Rodriguez-R LM, Phillippy AM, Konstantinidis KT, Aluru S. 2018. High throughput ANI analysis of 90K prokaryotic genomes reveals clear species boundaries. Nat Commun 9:5114.

41. Olm MR, Brown CT, Brooks B, Banfield JF. 2017. dRep: a tool for fast and accurate genomic comparisons that enables improved genome recovery from metagenomes through de-replication. ISME J 11:2864–2868.

42. Seemann T. 2017. barrnap: Bacterial ribosomal RNA predictor. Retrieved from 525.

43. Edgar RC. 2004. MUSCLE: multiple sequence alignment with high accuracy and high throughput. Nucleic Acids Res 32:1792–1797.

44. Minh BQ, Schmidt HA, Chernomor O, Schrempf D, Woodhams MD, von Haeseler A, Lanfear R. 2020. IQ-TREE 2: New Models and Efficient Methods for Phylogenetic Inference in the Genomic Era. Mol Biol Evol 37:1530–1534.

45. Letunic I, Bork P. 2021. Interactive Tree Of Life (iTOL) v5: an online tool for phylogenetic tree display and annotation. Nucleic Acids Res 49:W293–W296.

46. Camacho C, Coulouris G, Avagyan V, Ma N, Papadopoulos J, Bealer K, Madden TL. 2009. BLAST+: architecture and applications. BMC Bioinformatics 10:421.

47. Parks DH, Imelfort M, Skennerton CT, Hugenholtz P, Tyson GW. 2015. CheckM: assessing the quality of microbial genomes recovered from isolates, single cells, and metagenomes. Genome Res 25:1043–1055.

48. Getz EW, Tithi SS, Zhang L, Aylward FO. 2018. Parallel Evolution of Genome Streamlining and Cellular Bioenergetics across the Marine Radiation of a Bacterial Phylum. MBio 9.

49. Eren AM, Murat Eren A, Kiefl E, Shaiber A, Veseli I, Miller SE, Schechter MS, Fink I, Pan JN, Yousef M, Fogarty EC, Trigodet F, Watson AR, Esen ÖC, Moore RM, Clayssen Q, Lee MD, Kivenson V, Graham ED, Merrill BD, Karkman A, Blankenberg D, Eppley JM, Sjödin A, Scott JJ, Vázquez-Campos X, McKay LJ, McDaniel EA, Stevens SLR, Anderson RE, Fuessel J, Fernandez-Guerra A, Maignien L, Delmont TO, Willis AD. 2021. Community-led, integrated, reproducible multi-omics with anvi’o. Nature Microbiology https://doi.org/10.1038/s41564-020-00834-3.

50. Mistry J, Chuguransky S, Williams L, Qureshi M, Salazar GA, Sonnhammer ELL, Tosatto SCE, Paladin L, Raj S, Richardson LJ, Finn RD, Bateman A. 2021. Pfam: The protein families database in 2021. Nucleic Acids Res 49:D412–D419.

51. Tatusov RL, Galperin MY, Natale DA, Koonin EV. 2000. The COG database: a tool for genome-scale analysis of protein functions and evolution. Nucleic Acids Res 28:33–36.

52. Kanehisa M, Sato Y, Kawashima M, Furumichi M, Tanabe M. 2016. KEGG as a reference resource for gene and protein annotation. Nucleic Acids Res 44:D457–62.

53. Kanehisa M, Sato Y, Morishima K. 2016. BlastKOALA and GhostKOALA: KEGG Tools for Functional Characterization of Genome and Metagenome Sequences. J Mol Biol 428:726–731.

54. Shaiber A, Willis AD, Delmont TO, Roux S, Chen L-X, Schmid AC, Yousef M, Watson AR, Lolans K, Esen ÖC, Lee STM, Downey N, Morrison HG, Dewhirst FE, Mark Welch JL, Eren AM. 2020. Functional and genetic markers of niche partitioning among enigmatic members of the human oral microbiome. Genome Biol 21:292.

55. Eren AM, Esen ÖC, Quince C, Vineis JH, Morrison HG, Sogin ML, Delmont TO. 2015. Anvi’o: an advanced analysis and visualization platform for ‘omics data. PeerJ 3:e1319.

56. Buchfink B, Xie C, Huson DH. 2015. Fast and sensitive protein alignment using DIAMOND. Nat Methods 12:59–60.

57. Savoie ER, Lanclos VC, Henson MW, Cheng C, Getz EW, Barnes SJ, LaRowe DE, Rappé MS, Thrash JC. 2021. Ecophysiology of the Cosmopolitan OM252 Bacterioplankton (Gammaproteobacteria). mSystems e0027621.

58. Capella-Gutiérrez S, Silla-Martínez JM, Gabaldón T. 2009. trimAl: a tool for automated alignment trimming in large-scale phylogenetic analyses. Bioinformatics 25:1972–1973.

59. Ballesteros JA, Hormiga G. 2016. A New Orthology Assessment Method for Phylogenomic Data: Unrooted Phylogenetic Orthology. Mol Biol Evol 33:2117–2134.

60. Man D, Wang W, Sabehi G, Aravind L, Post AF, Massana R, Spudich EN, Spudich JL, Béjà O. 2003. Diversification and spectral tuning in marine proteorhodopsins. EMBO J 22:1725–1731.

61. Pesant S, Not F, Picheral M, Kandels-Lewis S, Le Bescot N, Gorsky G, Iudicone D, Karsenti E, Speich S, Troublé R, Dimier C, Searson S, Tara Oceans Consortium Coordinators. 2015. Open science resources for the discovery and analysis of Tara Oceans data. Sci Data 2:150023.

62. Alneberg J, Sundh J, Bennke C, Beier S, Lundin D, Hugerth LW, Pinhassi J, Kisand V, Riemann L, Jürgens K, Labrenz M, Andersson AF. 2018. BARM and BalticMicrobeDB, a reference metagenome and interface to meta-omic data for the Baltic Sea. Sci Data 5:180146.

63. Biller SJ, Berube PM, Dooley K, Williams M, Satinsky BM, Hackl T, Hogle SL, Coe A, Bergauer K, Bouman HA, Browning TJ, De Corte D, Hassler C, Hulston D, Jacquot JE, Maas EW, Reinthaler T, Sintes E, Yokokawa T, Chisholm SW. 2018. Marine microbial metagenomes sampled across space and time. Sci Data 5:180176.

64. Acinas SG, Sánchez P, Salazar G, Cornejo-Castillo FM, Sebastián M, Logares R, Sunagawa S, Hingamp P, Ogata H, Lima-Mendez G, Roux S, González JM, Arrieta JM, Alam IS, Kamau A, Bowler C, Raes J, Pesant S, Bork P, Agustí S, Gojobori T, Bajic V, Vaqué D, Sullivan MB, Pedrós-Alió C, Massana R, Duarte CM, Gasol JM. Metabolic Architecture of the Deep Ocean Microbiome https://doi.org/10.1101/635680.

65. Ahmed MA, Lim SJ, Campbell BJ. 2021. Metagenomes, Metatranscriptomes, and Metagenome-Assembled Genomes from Chesapeake and Delaware Bay (USA) Water Samples. Microbiol Resour Announc 10:e0026221.

66. Rasmussen AN, Francis CA. 2022. Genome-Resolved Metagenomic Insights into Massive Seasonal Ammonia-Oxidizing Archaea Blooms in San Francisco Bay. mSystems 7:e0127021.

67. Mende DR, Bryant JA, Aylward FO, Eppley JM, Nielsen T, Karl DM, DeLong EF. 2017. Environmental drivers of a microbial genomic transition zone in the ocean’s interior. Nature Microbiology https://doi.org/10.1038/s41564-017-0008-3.

68. Fortunato CS, Crump BC. 2015. Microbial Gene Abundance and Expression Patterns across a River to Ocean Salinity Gradient. PLoS One 10:e0140578.

69. Acinas SG, Sánchez P, Salazar G, Cornejo-Castillo FM, Sebastián M, Logares R, Royo-Llonch M, Paoli L, Sunagawa S, Hingamp P, Ogata H, Lima-Mendez G, Roux S, González JM, Arrieta JM, Alam IS, Kamau A, Bowler C, Raes J, Pesant S, Bork P, Agustí S, Gojobori T, Vaqué D, Sullivan MB, Pedrós-Alió C, Massana R, Duarte CM, Gasol JM. 2021. Deep ocean metagenomes provide insight into the metabolic architecture of bathypelagic microbial communities. Communications Biology https://doi.org/10.1038/s42003-021-02112-2.

70. Kojima CY, Getz EW, Thrash JC. 2022. RRAP: RPKM Recruitment Analysis Pipeline. Microbiol Resour Announc e0064422.

71. Langmead B. 2013. Bowtie2 manual. Dosegljivo: https://githubcom/BenLangmead/bowtie2/blob/master/MANUAL,[Dostopano: 16 7 2018].

72. Li H, Handsaker B, Wysoker A, Fennell T, Ruan J, Homer N, Marth G, Abecasis G, Durbin R, 1000 Genome Project Data Processing Subgroup. 2009. The Sequence Alignment/Map format and SAMtools. Bioinformatics 25:2078–2079.

73. Schlitzer R. 2002. Interactive analysis and visualization of geoscience data with Ocean Data View. Comput Geosci 28:1211–1218.

74. Thrash JC, Weckhorst JL, Pitre DM. 2015. Cultivating Fastidious Microbes, p. 57–78. In McGenity, TJ, Timmis, KN, Nogales, B (eds.), Springer Protocols Handbooks. Springer Berlin Heidelberg, Berlin, Heidelberg.

75. Yarza P, Yilmaz P, Pruesse E, Glöckner FO, Ludwig W, Schleifer K-H, Whitman WB, Euzéby J, Amann R, Rosselló-Móra R. 2014. Uniting the classification of cultured and uncultured bacteria and archaea using 16S rRNA gene sequences. Nat Rev Microbiol 12:635–645.

76. R Core Team. 2021. R: A Language and Environment for Statistical Computing. R Foundation for Statistical Computing, Vienna, Austria.

77. 隆伊藤. 1959. The Venice system for the classification of marine waters according to salinity : Symposium on the classification of brackish waters, Venice, 8-14 April 1958. 陸水学雑誌 20:119–120.

78. Henson MW, Lanclos VC, Faircloth BC, Thrash JC. 2018. Cultivation and genomics of the first freshwater SAR11 (LD12) isolate. ISME J 12:1846–1860.

79. Tolbert NE, Zill LP. 1956. Excretion of glycolic acid by algae during photosynthesis. J Biol Chem 222:895–906.

80. Wright RT, Shah NM. 1977. The trophic role of glycolic acid in coastal seawater. II. Seasonal changes in concentration and heterotrophic use in Ipswich Bay, Massachusetts, USA. Mar Biol 43:257–263.

81. Haro-Moreno JM, López-Pérez M, Alekseev A, Podoliak E, Kovalev K, Gordeliy V, Stepanauskas R, Rodriguez-Valera F. 2023. Flotillin-Associated rhodopsin (FArhodopsin), a widespread paralog of proteorhodopsin in aquatic bacteria with streamlined genomes. bioRxiv.

82. Thrash JC, Boyd A, Huggett MJ, Grote J, Carini P, Yoder RJ, Robbertse B, Spatafora JW, Rappé MS, Giovannoni SJ. 2011. Phylogenomic evidence for a common ancestor of mitochondria and the SAR11 clade. Scientific Reports https://doi.org/10.1038/srep00013.

83. Carlson CA, Morris R, Parsons R, Treusch AH, Giovannoni SJ, Vergin K. 2009. Seasonal dynamics of SAR11 populations in the euphotic and mesopelagic zones of the northwestern Sargasso Sea. ISME J 3:283–295.

84. Treusch AH, Vergin KL, Finlay LA, Donatz MG, Burton RM, Carlson CA, Giovannoni SJ. 2009. Seasonality and vertical structure of microbial communities in an ocean gyre. ISME J 3:1148–1163.

85. Schattenhofer M, Fuchs BM, Amann R, Zubkov MV, Tarran GA, Pernthaler J. 2009. Latitudinal distribution of prokaryotic picoplankton populations in the Atlantic Ocean. Environ Microbiol 11:2078–2093.

86. Alonso-Sáez L, Balagué V, Sà EL, Sánchez O, González JM, Pinhassi J, Massana R, Pernthaler J, Pedrós-Alió C, Gasol JM. 2007. Seasonality in bacterial diversity in north-west Mediterranean coastal waters: assessment through clone libraries, fingerprinting and FISH. FEMS Microbiol Ecol 60:98–112.

87. Mou X, Vila-Costa M, Sun S, Zhao W, Sharma S, Moran MA. 2011. Metatranscriptomic signature of exogenous polyamine utilization by coastal bacterioplankton. Environ Microbiol Rep 3:798–806.

88. Lu X, Sun S, Hollibaugh JT, Mou X. 2015. Identification of polyamine-responsive bacterioplankton taxa in South Atlantic Bight. Environ Microbiol Rep 7:831–838.

89. Noell SE, Barrell GE, Suffridge C, Morré J, Gable KP, Graff JR, VerWey BJ, Hellweger FL, Giovannoni SJ. 2021. SAR11 Cells Rely on Enzyme Multifunctionality To Metabolize a Range of Polyamine Compounds. MBio 12:e0109121.

90. Igarashi K, Kashiwagi K. 2010. Characteristics of cellular polyamine transport in prokaryotes and eukaryotes. Plant Physiol Biochem 48:506–512.

91. Hogle SL, Thrash JC, Dupont CL, Barbeau KA. 2016. Trace Metal Acquisition by Marine Heterotrophic Bacterioplankton with Contrasting Trophic Strategies. Appl Environ Microbiol 82:1613–1624.

92. Kim S, Kang I, Lee J-W, Jeon CO, Giovannoni SJ, Cho J-C. 2021. Heme auxotrophy in abundant aquatic microbial lineages. Proc Natl Acad Sci U S A 118.

93. Buessecker S, Palmer M, Lai D, Dimapilis J, Mayali X, Mosier D, Jiao J-Y, Colman DR, Keller LM, St John E, Miranda M, Gonzalez C, Gonzalez L, Sam C, Villa C, Zhuo M, Bodman N, Robles F, Boyd ES, Cox AD, St Clair B, Hua Z-S, Li W-J, Reysenbach A-L, Stott MB, Weber PK, Pett-Ridge J, Dekas AE, Hedlund BP, Dodsworth JA. 2022. An essential role for tungsten in the ecology and evolution of a previously uncultivated lineage of anaerobic, thermophilic Archaea. Nat Commun 13:3773.

94. Kletzin A, Adams MW. 1996. Tungsten in biological systems. FEMS Microbiol Rev 18:5–63.

95. Coimbra C, Farias P, Branco R, Morais PV. 2017. Tungsten accumulation by highly tolerant marine hydrothermal Sulfitobacter dubius strains carrying a tupBCA cluster. Syst Appl Microbiol 40:388–395.

96. Ferry JG. 1990. Formate dehydrogenase. FEMS Microbiol Rev 7:377–382.

97. Carini P, Campbell EO, Morré J, Sañudo-Wilhelmy SA, Thrash JC, Bennett SE, Temperton B, Begley T, Giovannoni SJ. 2014. Discovery of a SAR11 growth requirement for thiamin’s pyrimidine precursor and its distribution in the Sargasso Sea. ISME J 8:1727–1738.

98. Cheng C, Thrash JC. 2021. sparse-growth-curve: a Computational Pipeline for Parsing Cellular Growth Curves with Low Temporal Resolution. Microbiol Resour Announc 10.

